## Supplemental Figures and summary of Supplemental Tables for "Global genome diversity of the *Leishmania donovani* complex"

July 19, 2019

### 1 Figures

#### 1.1 Evolution in the *L. donovani* complex

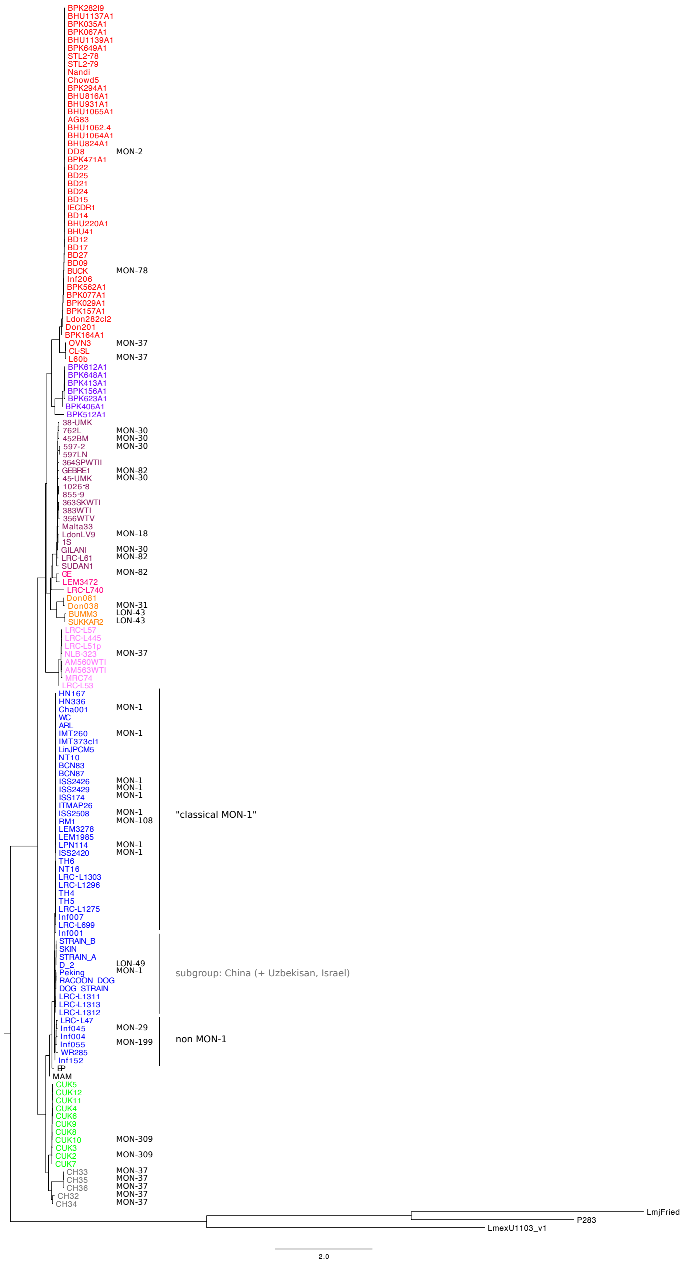

Figure 1

Figure 1: Phylogenetic reconstruction of all 151 samples of the *L. donovani* complex. The phylogeny was calculated with neighbour joining based on Nei's distances including isolates of *L. mexicana* (U1103.v1), *L. tropica* (P283) and *L. major* (LmjFried) as outgroups. Group colours are identical to those used throughout the manuscript. Phylogenetic grouping as suggested by Multilocus Enzyme Electrophoresis are indicated in black (see Table S1 for references).

#### 1.2 Aneuploidy

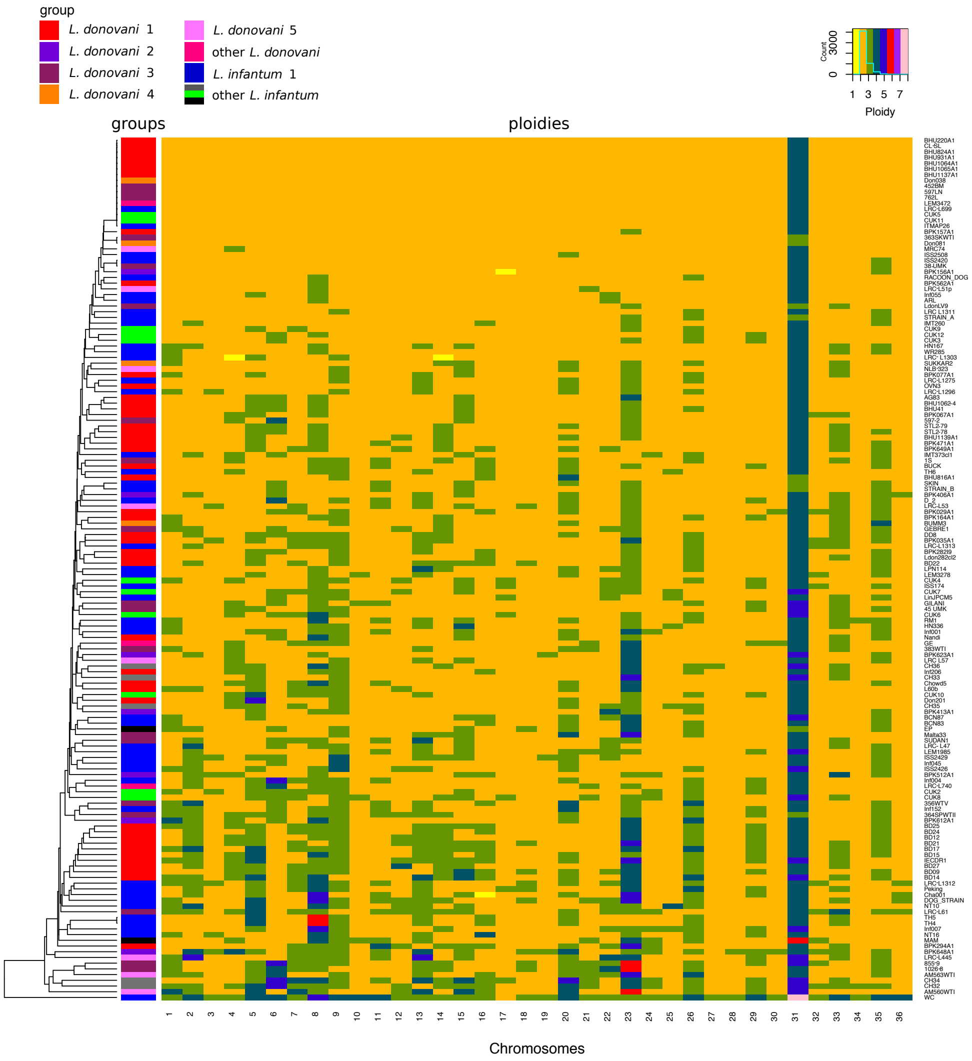

Figure 2: Aneuploidy patterns across all 151 samples. The heatmap displays the ploidies of the individual chromosomes and samples. Samples, displayed in different rows, are ordered by average linkage clustering and chromosomes are shown in different columns. Different ploidies are indicated by the colours in the heatmap (legend: upper right corner). The column on the left indicates the different phylogenetic groups (legend: upper left corner).

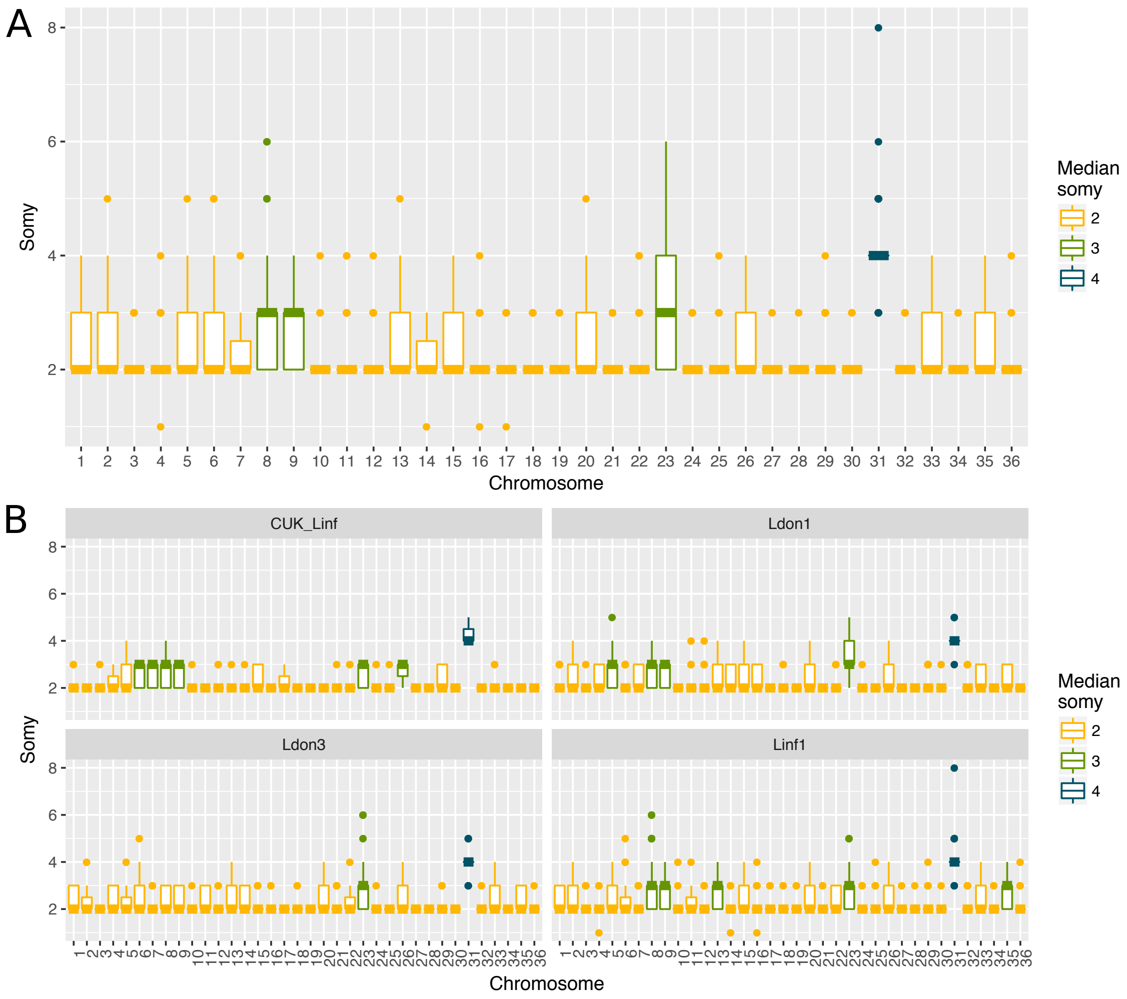

Figure 3: Aneuploidy distributions for the different chromosomes. A) The boxplots show the distributions of all observed ploidies across all 151 samples for each chromosome. B) Ploidy distributions are shows across the four largest (sub-)groups (>=9 samples per group) identified in the data set.

#### 1.3 Heterozygosity

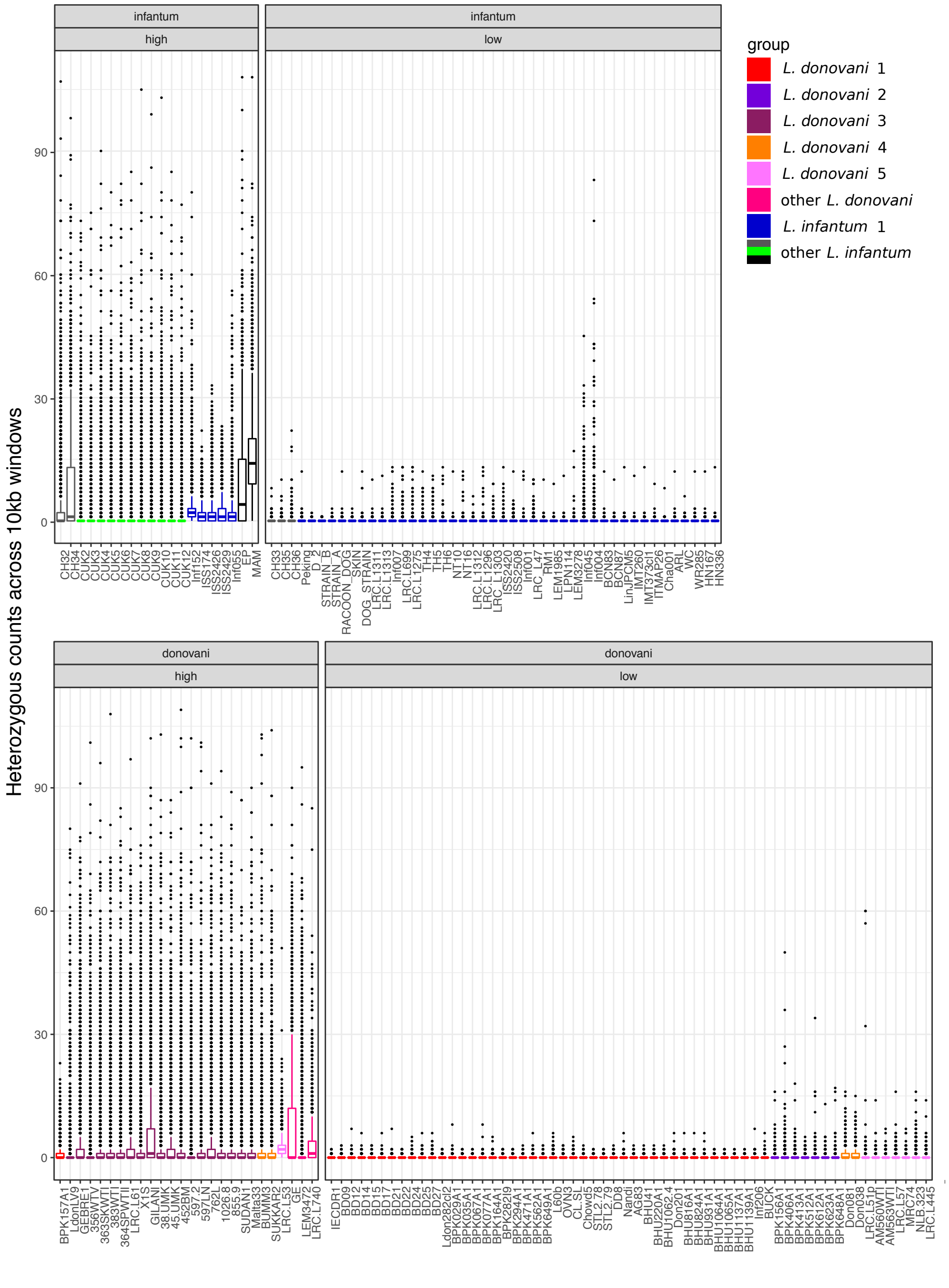

Figure 4

Figure 4: Distribution of heterozygous sites across the genome. Boxplots show the distribution of heterozygous counts for 10 kb windows across the genome for each sample. Upper panels show sample categorisation by species and genome-wide ”high” versus ”low” heterozygosity samples (i.e. above or below a genome-wide heterozygosity of 0.004). Groups are indicated by the different colours used throughout this study.

#### 1.4 Genomic signatures of hybridisation

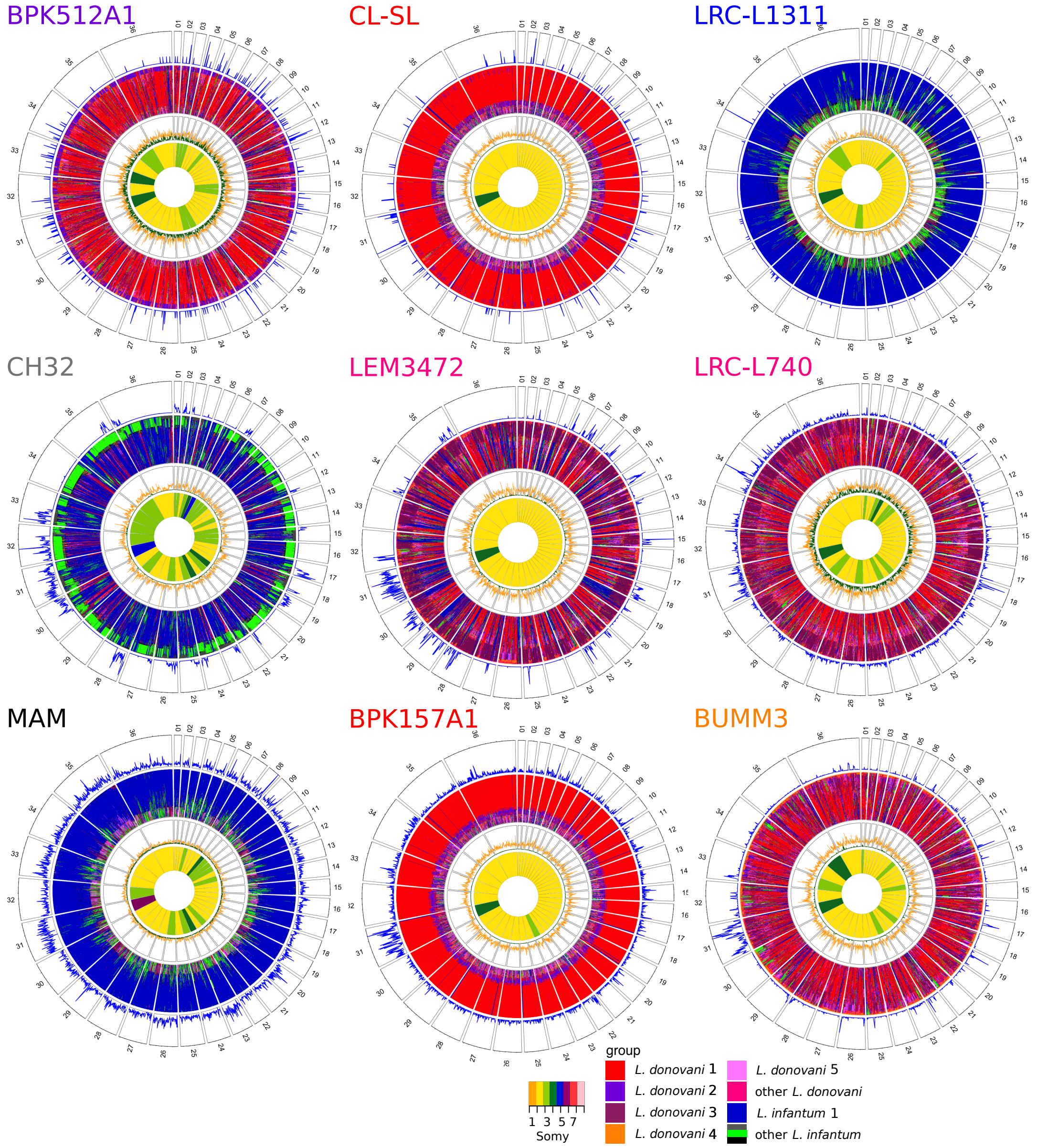

(a) Each circos plot shows four different genomic features of the isolate named in each top left corner. In the four different rings, pies correspond to the different chromosomes labeled by the chromosome number. The three outer rings show a window based analysis for a window size of 10 kb. Starting from the outer ring, they show: 1. Heterozygosity (number of heterozygous sites; range from 0 to 5 (BPK512A1), 5 (CL-SL), 10 (LRC-L1311), 107 (CH32), 95 (LEM3472), 85 (LRC-L740), 108 (MAM), 23 (BPK157A1) and 103 (BUMM3) heterozygous sites per 10 kb, respectively), 2. A heatmap coloured by groups of the 60 genetically closest isolates (based on Nei's D), starting with the closest sample at the outer margin and the $60^{th}$ furthest isolate at the inner margin, 3. Nei's D to the closest isolate (green) and the $60^{th}$ closest sample (orange), range from 0 to 1. The innermost circle shows the colour-coded somy.

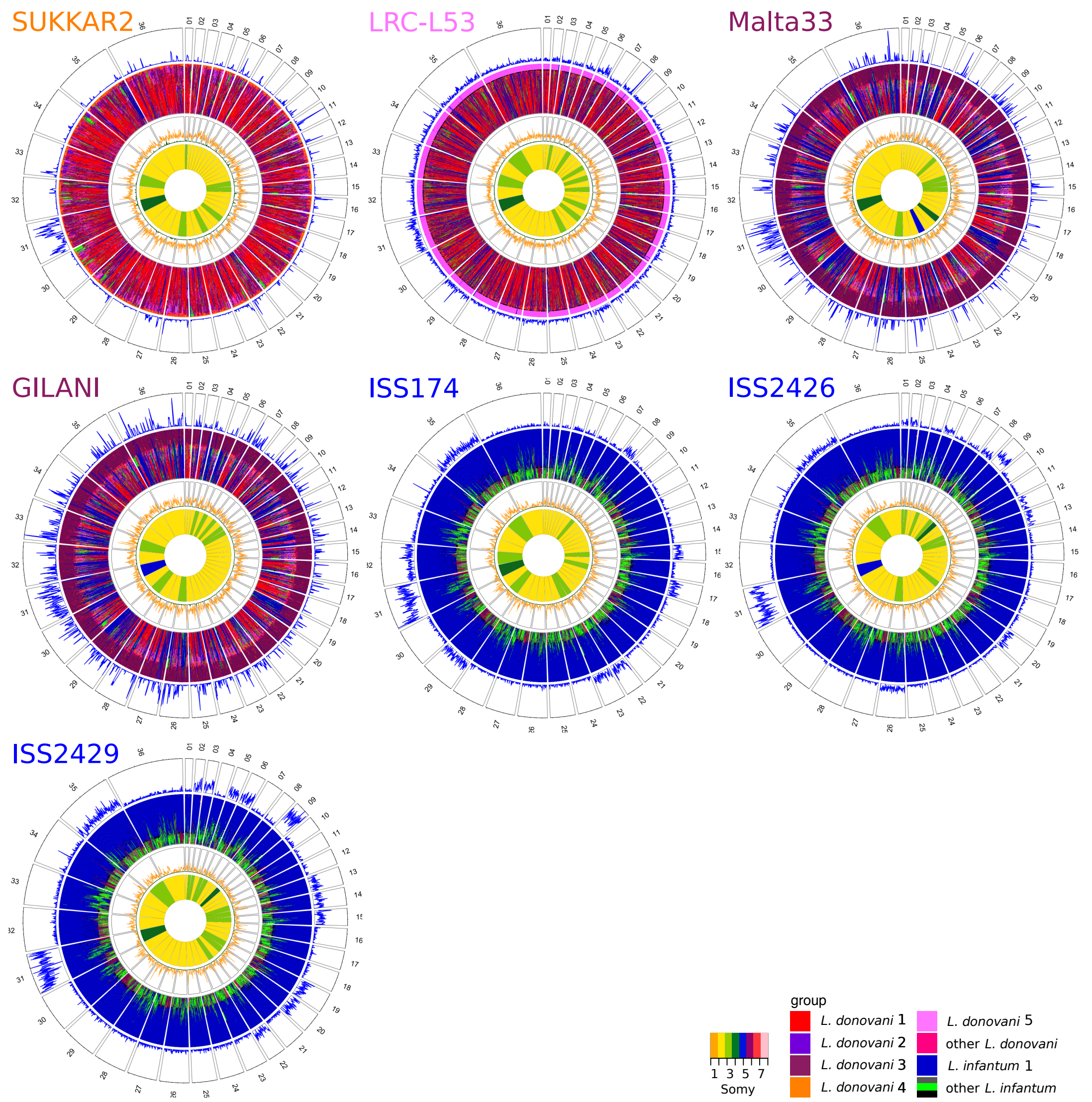

(b) Each circos plot shows four different genomic features of the isolate named in each top left corner. In the four different rings, pies correspond to the different chromosomes labeled by the chromosome number. The three outer rings show a window based analysis for a window size of 10 kb. Starting from the outer ring, they show: 1. Heterozygosity (number of heterozygous sites; range from 0 to 104 (SUKKAR2), 51 (LRC-L53), 90 (Malta33), 102 (GILANI), 22 (ISS174), 33 (ISS2426) and 23 (ISS2429) heterozygous sites per 10 kb, respectively), 2. A heatmap coloured by groups of the 60 genetically closest isolates (based on Nei's D), starting with the closest sample at the outer margin and the $60^{th}$ furthest isolate at the inner margin, 3. Nei's D to the closest isolate (green) and the $60^{th}$ closest sample (orange), range from 0 to 1. The innermost circle shows the colour-coded somy.

Figure 5: Window based analysis of relatedness for a subset of samples.

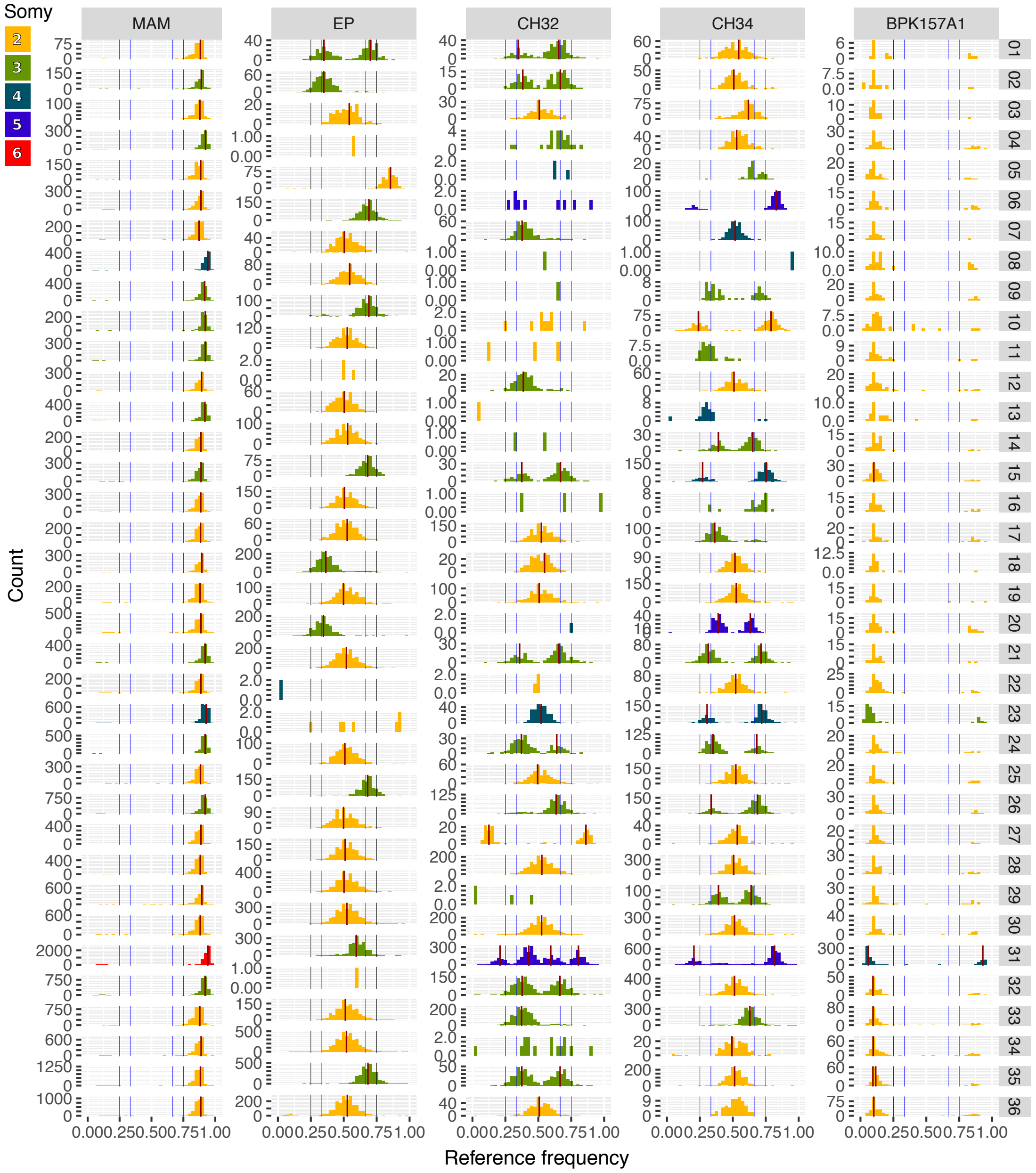
Somy
2
3
4
5
6
MAM
EP
CH32
CH34
BPK157A1
Count
Reference frequency
0.000.250.500.751.00

(a)

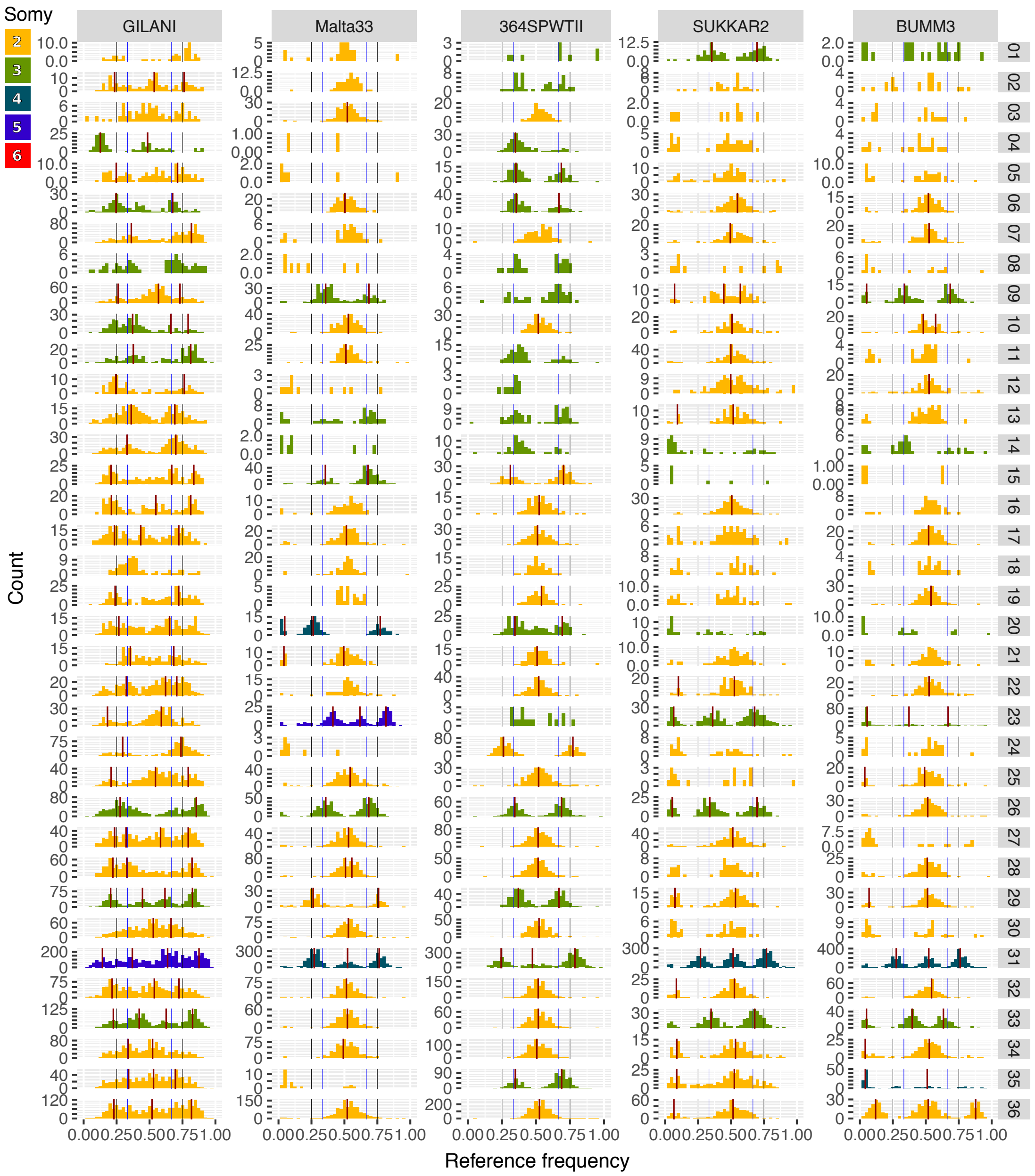
Somy
2
3
4
5
6
GILANI
Malta33
364SPWTII
SUKKAR2
BUMM3
Count
Reference frequency
0.000.250.500.751.00
01 02 03 04 05 06 07 08 09 10 11 12 13 14 15 16 17 18 19 20 21 22 23 24 25 26 27 28 29 30 31 32 33 34 35 36

(b)

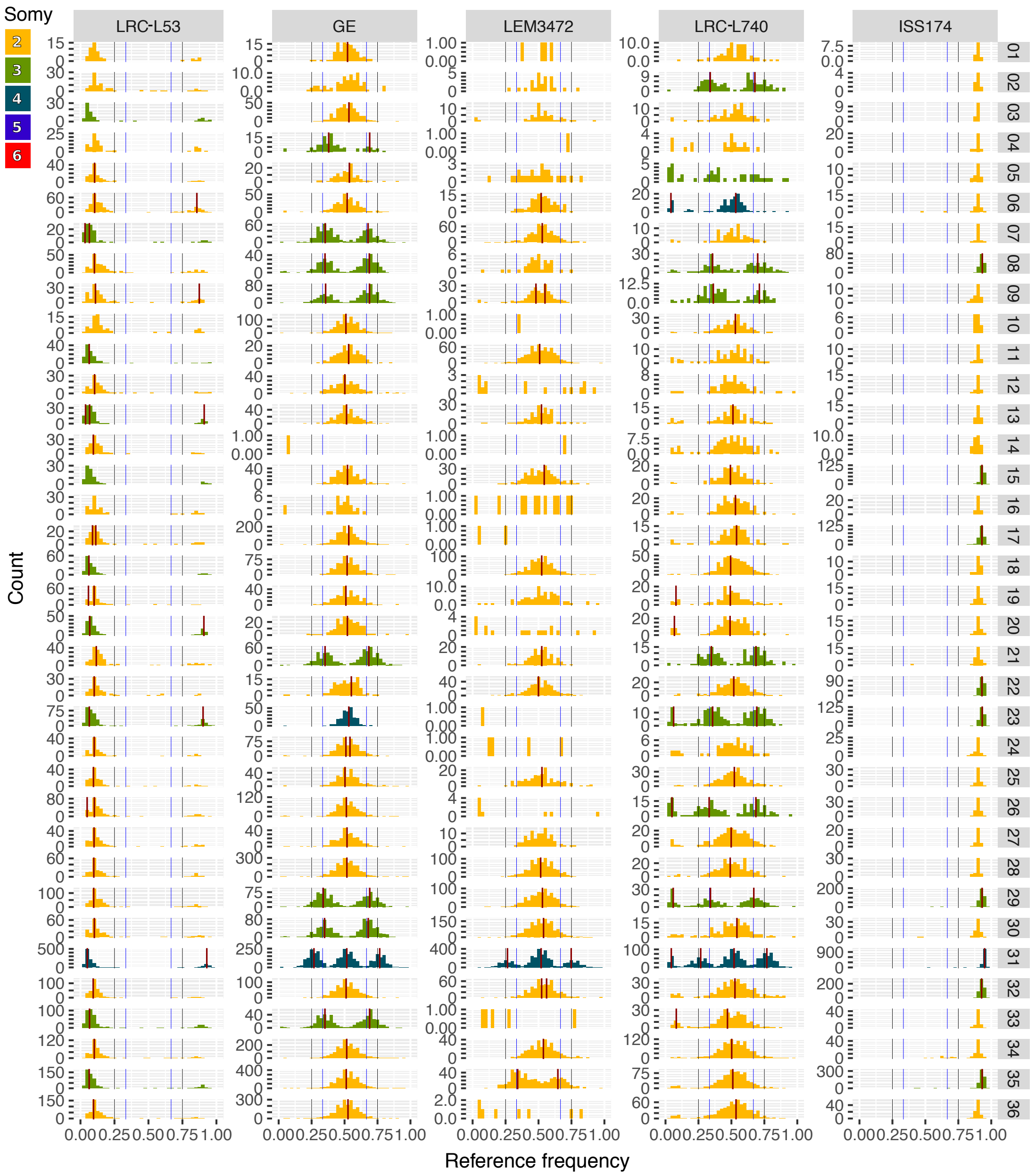
Somy
2
3
4
5
6
LRC-L53
GE
LEM3472
LRC-L740
ISS174
Count
Reference frequency
0.00 0.25 0.50 0.75 1.00

(c)

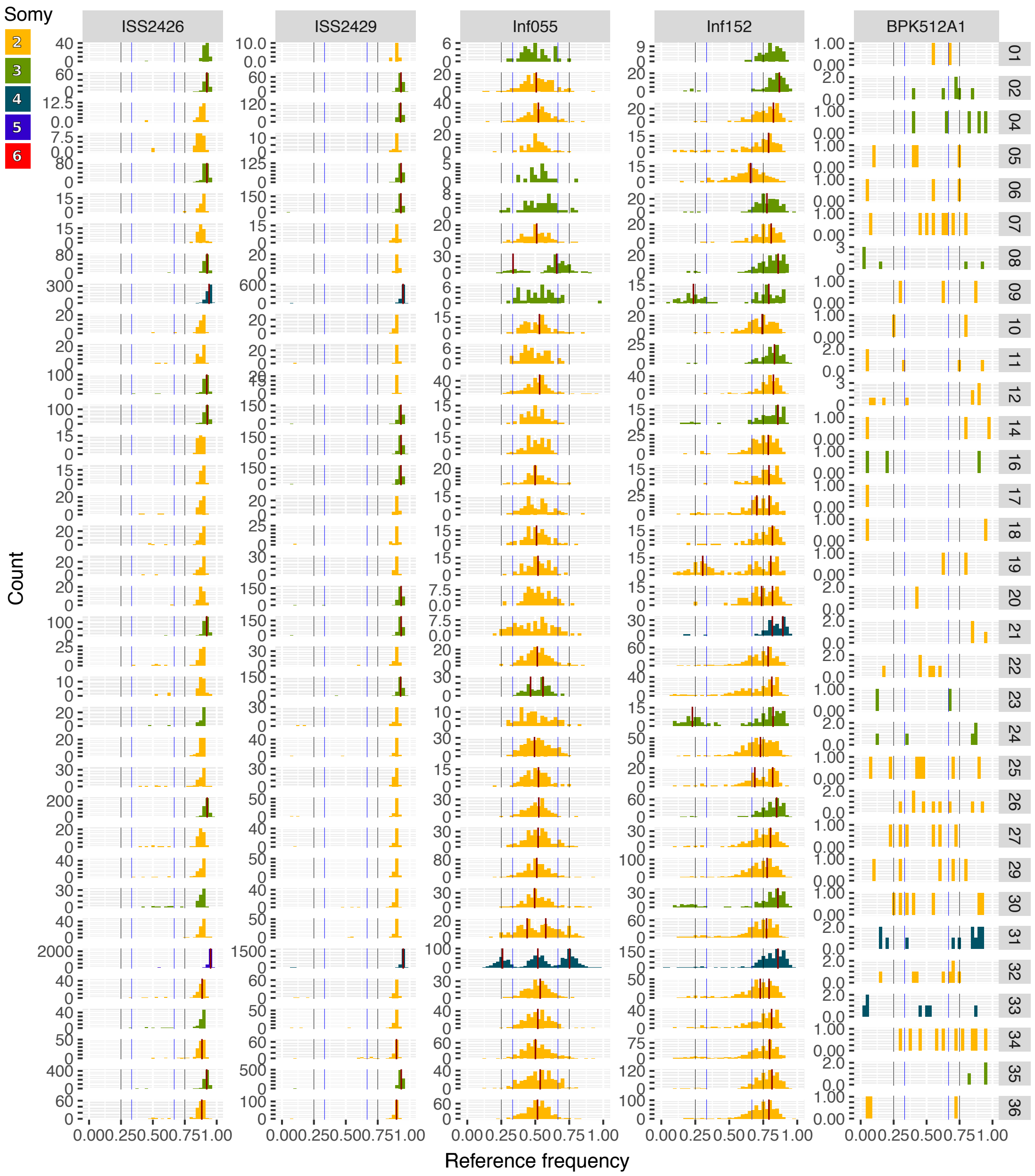

(d)

Figure 6: Allele frequency distributions by isolate. Histograms of allele frequency distributions are shown per chromosome for different isolates indicated at the top of each plot. Allele frequencies of 1/3 & 2/3 and 1/4 & 3/4 are visualised by blue and black vertical lines, respectively. Red vertical lines indicate estimated peaks of the distribution, which are only shown for distributions with at least 100 SNPs (see Material & Methods). Colours of the distributions indicate the estimated somy based on chromosomal coverage.

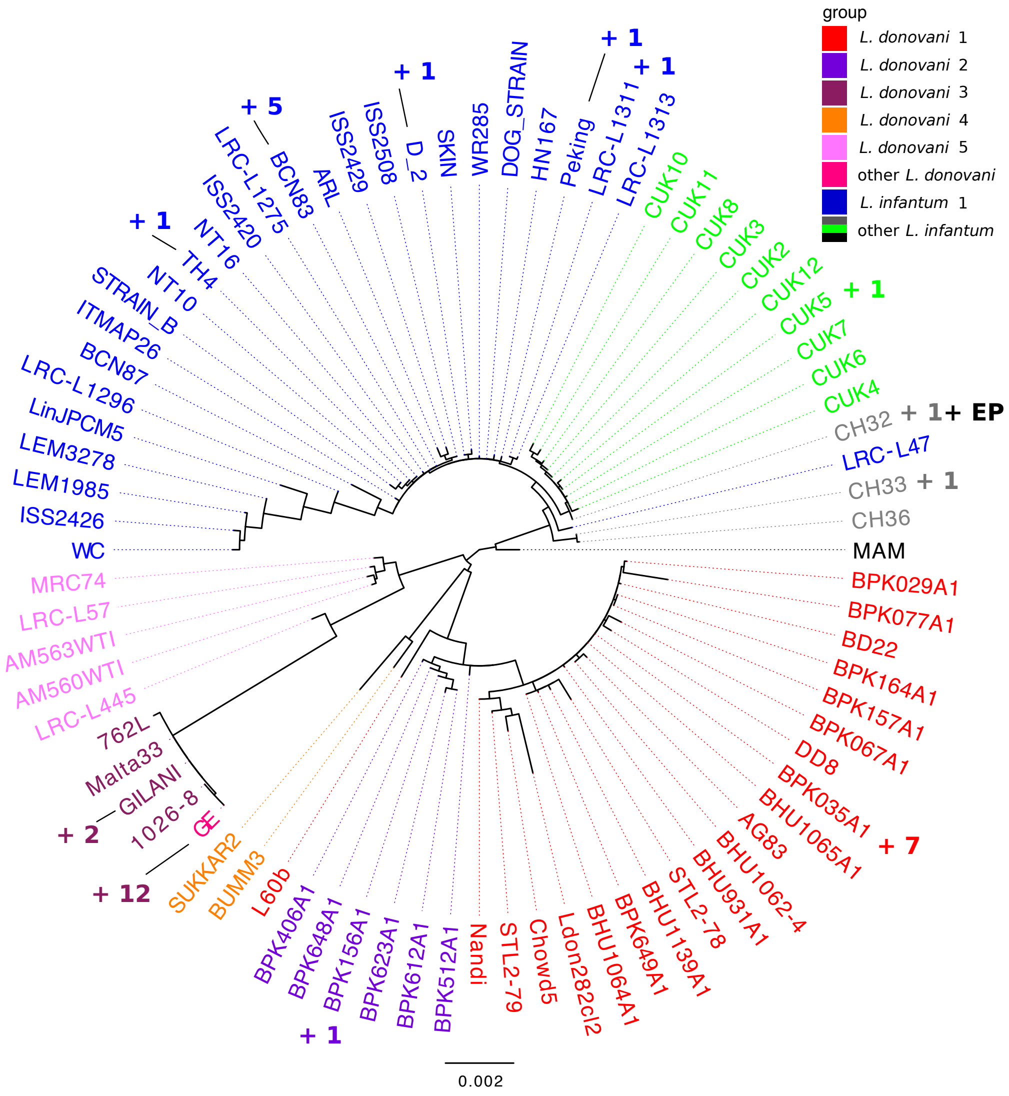

Figure 7: Phylogenetic tree based on the maxicircle DNA. Unrooted tree constructed using RaxML (see Material & Methods). Only a part of the samples is shown in the tree as the library preparation of some samples removed the kDNA and some samples have identical sequences to the ones shown and were therefore removed for tree estimation. The samples with identical sequences are listed in table S2 and their numbers are indicated near the respective tree leaves. Group identity of each isolate is indicated by the different colours. Phylogenetic reconstruction was repeated with four outgroup isolates of *L. mexicana* (U1103.v1), *L. tropica* (P283) and *L. amazonensis* (two samples) all of those four samples used were clearly different to the samples of our study.

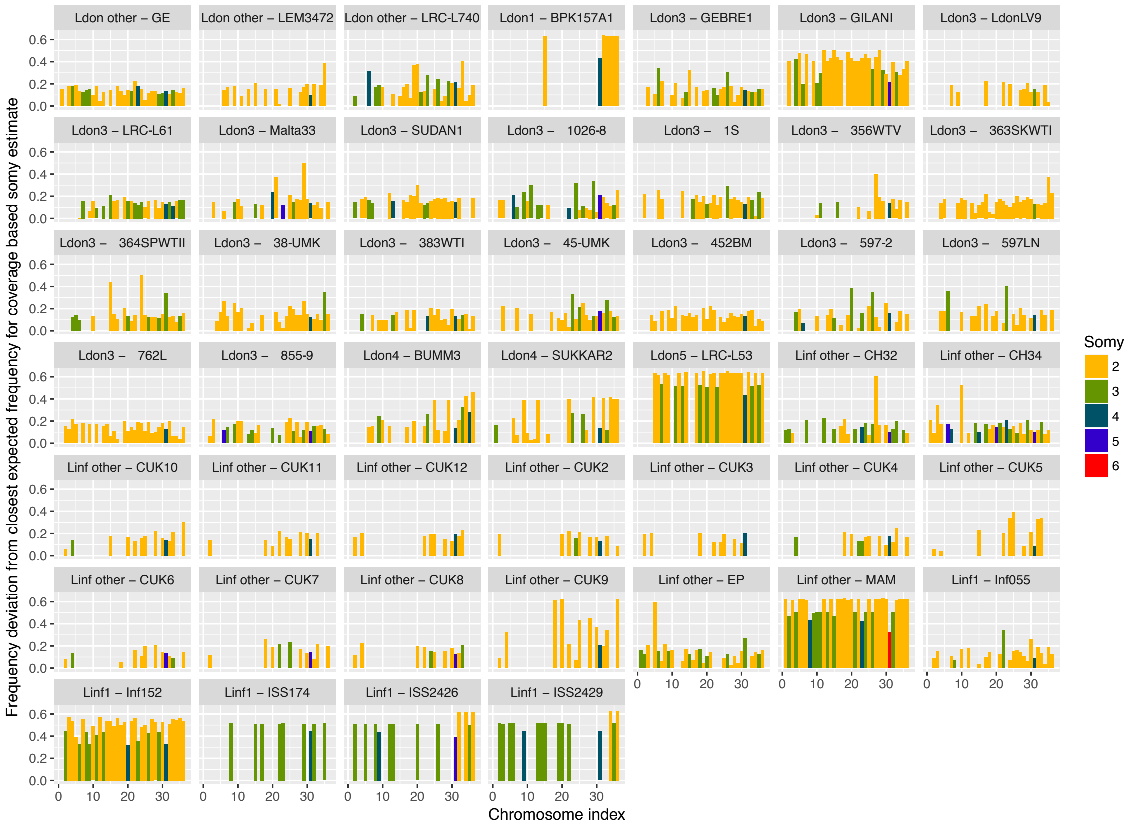

Figure 8: Somy evaluation based on allele frequency profiles. Somies for the different isolates and chromosomes were calculated based on relative chromosome wide coverages within a sample. For heterozygous samples and chromosomes with at least 100 SNPs, these estimates were evaluated based on the expected frequency distributions (see supplemental Material & Methods "Somy evaluation based on allele frequency profiles"). Errors above 0.2 suggest deviances in somy estimates larger than expected by sampling error. In a few cases, the frequency profiles clearly suggest another somy, than estimated by chromosomal coverage.

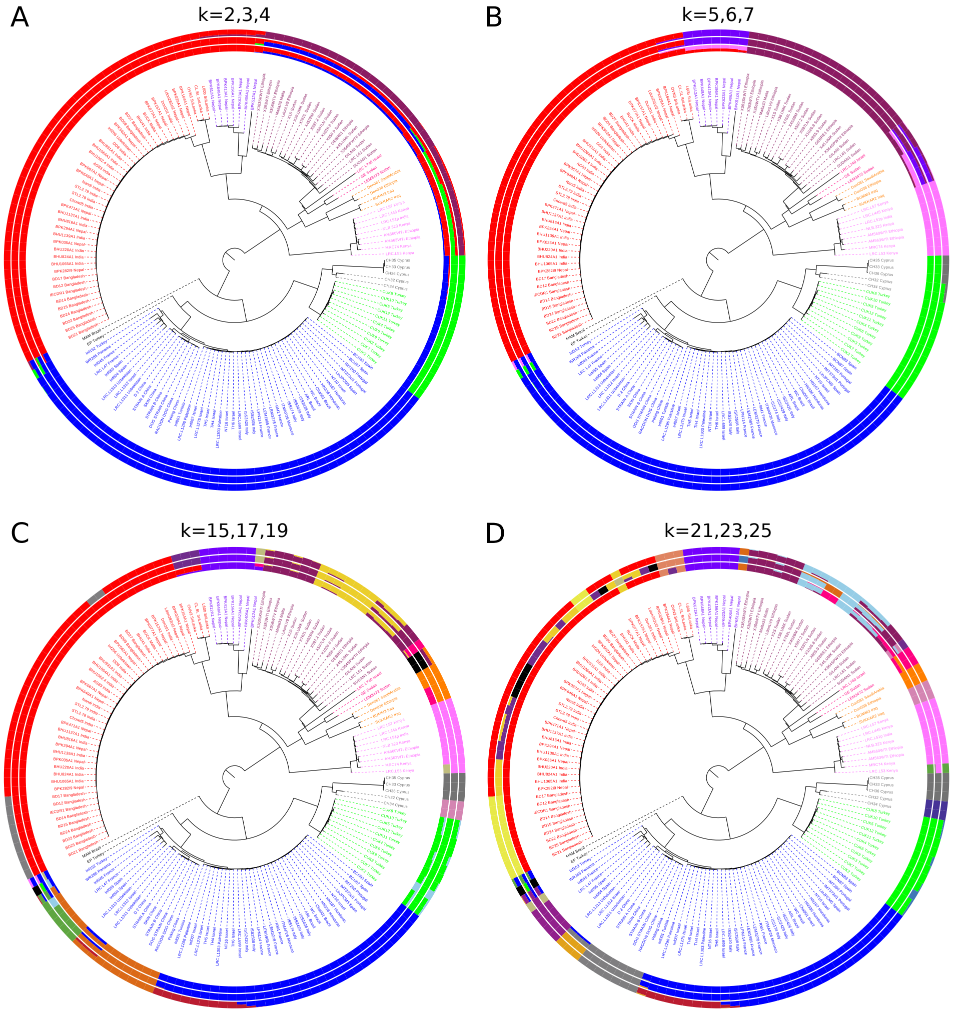

Figure 9: Sample phylogeny and admixture analysis across a range of k values. The phylogenetic reconstruction is identical to the figure 1 but admixture results shown are for a range of different k values. Group colours in the phylogeny are identical to the ones used throughout this study and are matched for admixture results where possible.

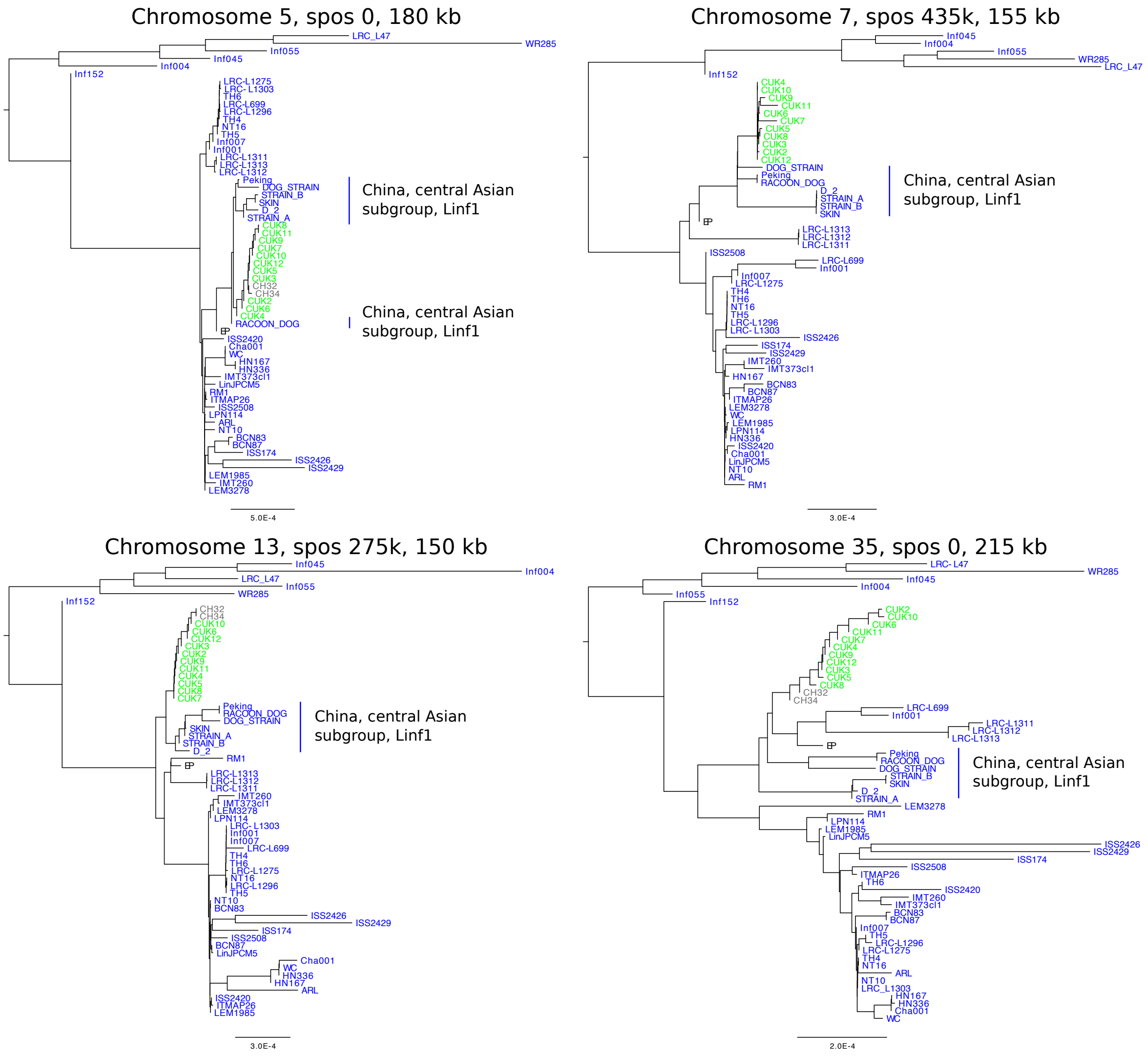
Chromosome 5, spos 0, 180 kb
China, central Asian subgroup, Linf1
China, central Asian subgroup, Linf1
5.0E-4
Chromosome 7, spos 435k, 155 kb
China, central Asian subgroup, Linf1
3.0E-4
Chromosome 13, spos 275k, 150 kb
China, central Asian subgroup, Linf1
3.0E-4
Chromosome 35, spos 0, 215 kb
China, central Asian subgroup, Linf1
2.0E-4

(a)

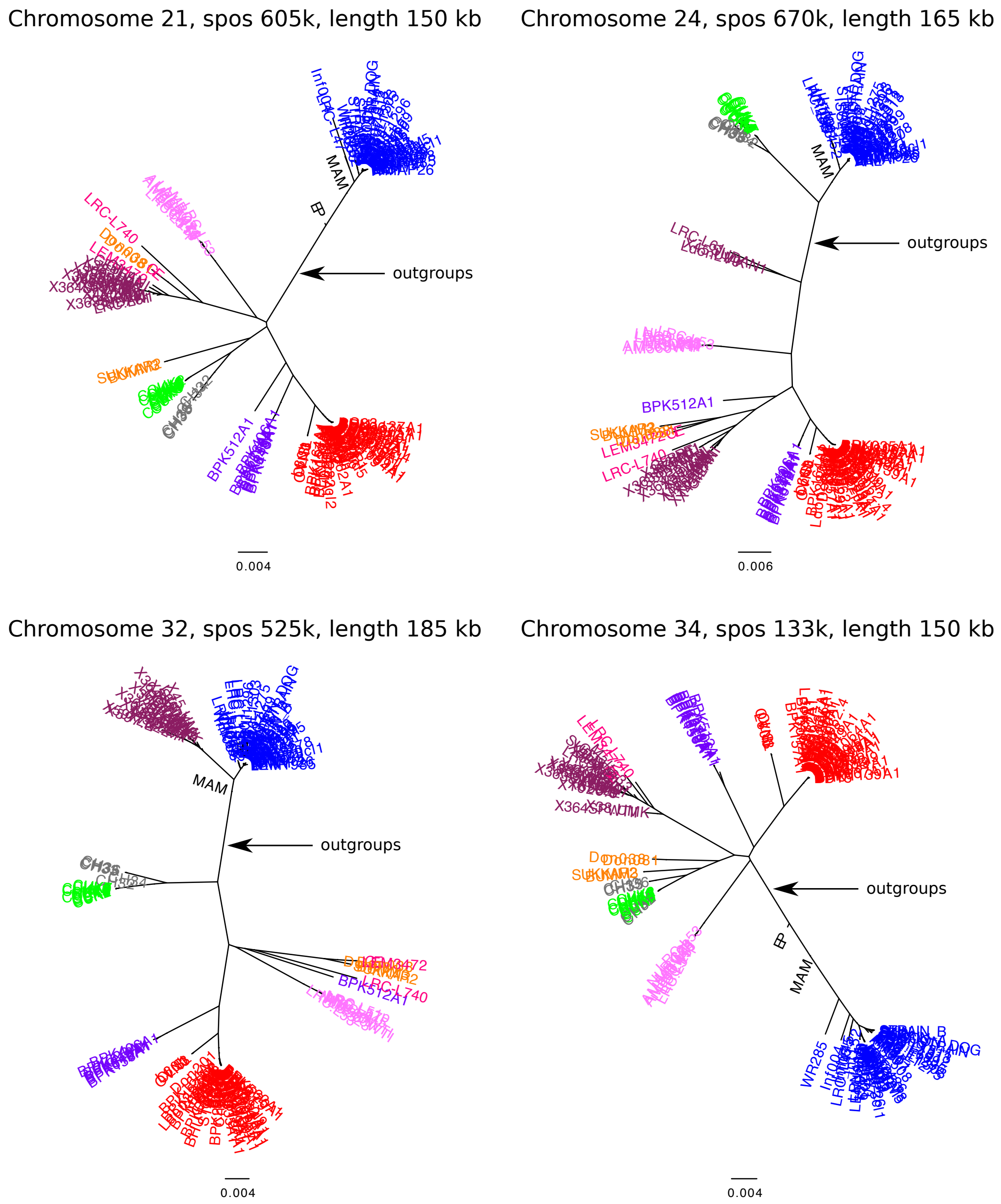
Chromosome 21, spos 605k, length 150 kb
Chromosome 24, spos 670k, length 165 kb
Chromosome 32, spos 525k, length 185 kb
Chromosome 34, spos 133k, length 150 kb
outgroups
MAM
BP
BPK512A1
LRC-L740
WR285
0.004
0.006

(b)

Figure 10: Putative parents of CUK samples. Phylogenetic trees were reconstructed using Nei's D and neighbour joining including all 151 samples and three outgroup samples, *L. mexicana* (U1103.v1), *L. tropica* (P283) (see Material & Methods). Trees were done for the four largest homozygous genome regions across the CUK genomes either almost devoid of differences (a) or with increased fixed differences (b) to the JPCM5 reference (see Material & Methods, "Haplotype-based analysis of hybridisation in CUK isolates"). (a) Phylogenetic trees using genomic regions putatively from a JPCM5-like parent in the CUK samples. For a better resolution only the subtree for the Linf1 clade is shown. (b) Phylogenetic trees using genomic regions with increased numbers of fixed differences to the JPCM5 reference in the CUK samples. For a better resolution the the position of the out-group branch is only indicated.

#### 1.5 Isolates with genetically distinct (sub-)clones

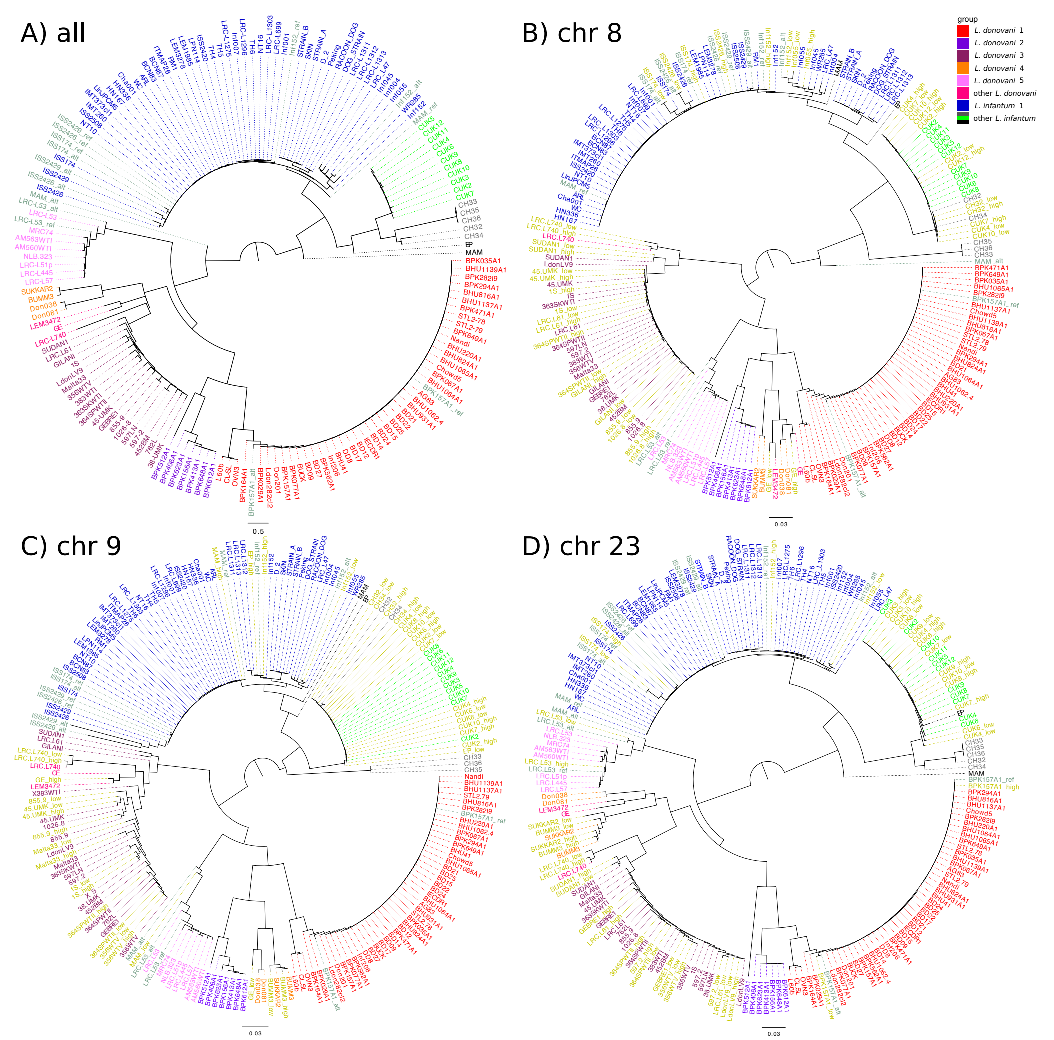

Figure 11: Sample phylogeny based on genomic SNP variation including phased samples with skewed allele frequency spectra. Phylogenetic reconstruction was equivalent to the reconstruction shown in figure 1. A-D) Samples with strongly skewed allele frequency spectra across all chromosomes were phased based on high versus low allele frequency variants and also included in the tree. Resulting haplotypes are coloured in greenish grey and labelled with ref (reference allele; reference *L. infantum* JPCM5) and alt (alternate allele) depending on the polarisation of the majority if SNPs in the respective haplotype. B-D) Individual chromosomes that were triploid were phased based on the allele frequency spectra. Resulting haplotypes are coloured in yellowish grey and labeled with "high", where the JPCM5 reference allele is at high ($\sim 2/3$) and the non-reference allele is at low ($\sim 1/3$) sample frequency. The haplotype resulting from the opposite scenario is labeled with "low". The remaining samples are coloured in their typical group colours.

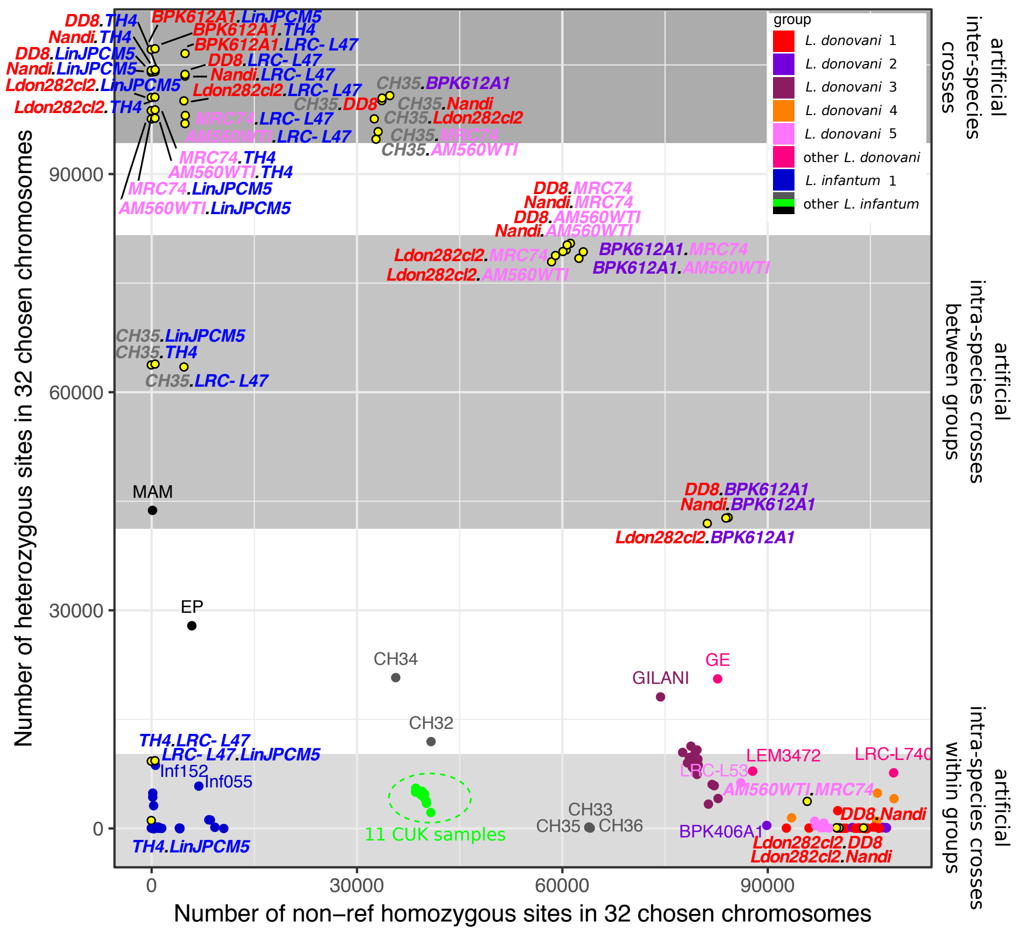

Figure 12: Heterozygosity of artificial F1 hybrids. Artificial F1 hybrids were generated by matching ten almost entirely homozygous samples across the phylogeny. Chromosomes 2, 8, 22 and 31 were excluded since they did not satisfy these criteria. The number of non-reference homozygous sites is plotted against the number of heterozygous sites. The plot includes real samples as well as artificial hybrids. Groups are indicated by colours. Artificial hybrids are displayed by yellow dots surrounded by black circles and names are written in bold italics indicating the names of the combined samples in their respective group colours separated by a black dot. Grey shaded areas indicate heterozygosity levels that are spanned by different levels of artificial hybrids including inter-species and intra-species hybrids between and within identified groups.

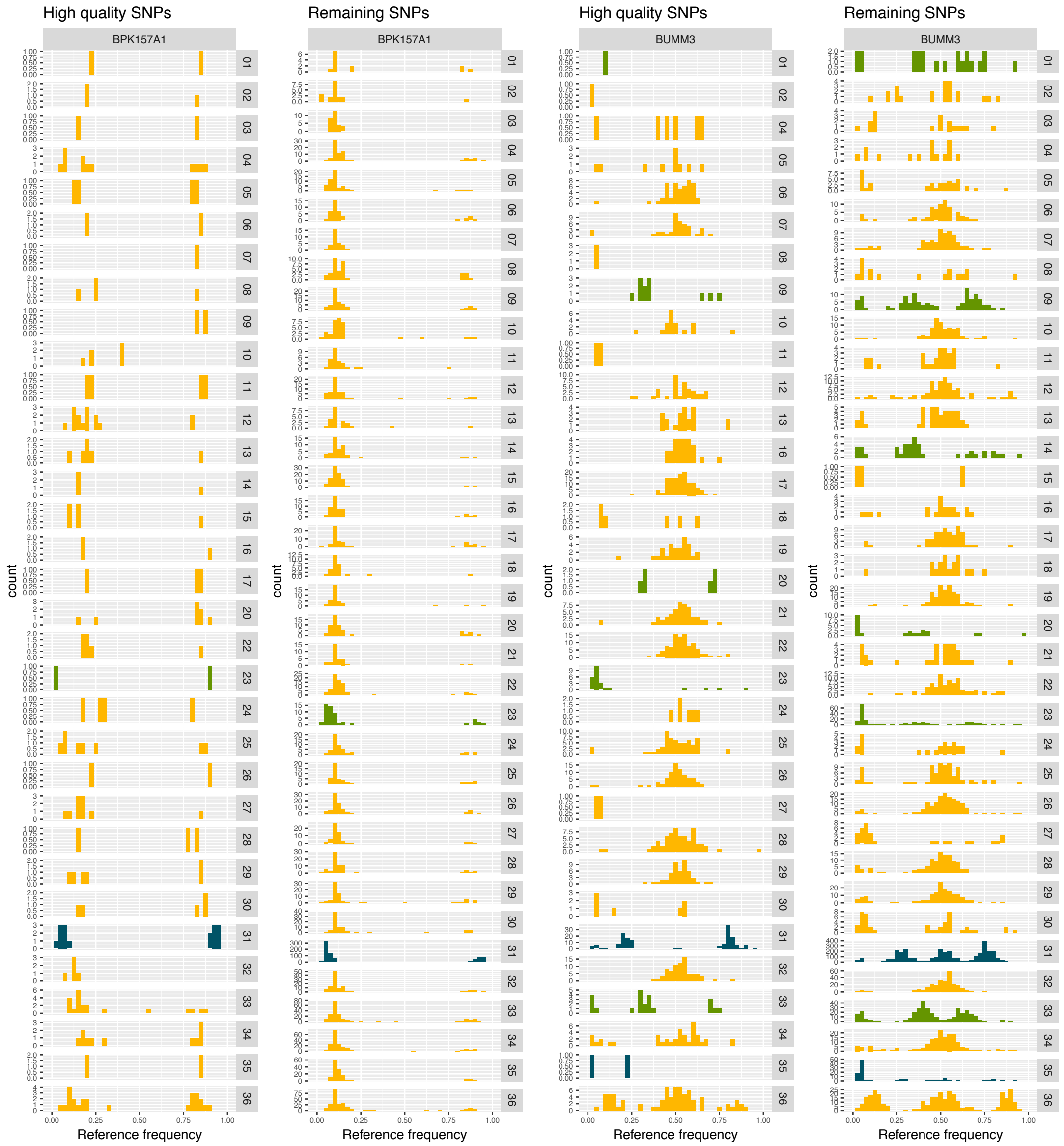
High quality SNPs
Remaining SNPs
High quality SNPs
Remaining SNPs
BPK157A1
BPK157A1
BUMM3
BUMM3
count
Reference frequency

(a)

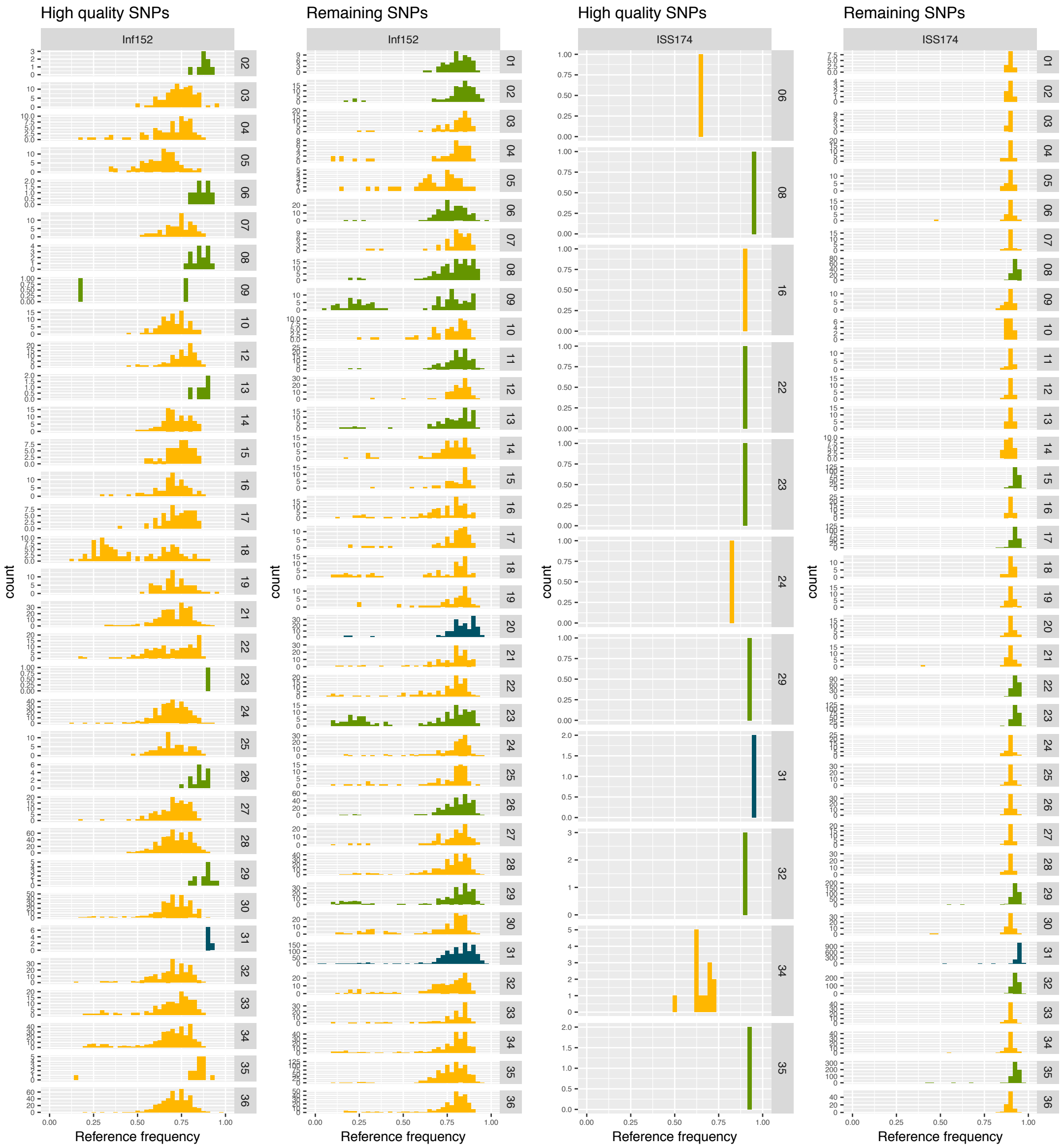
High quality SNPs
Inf152
Remaining SNPs
Inf152
High quality SNPs
ISS174
Remaining SNPs
ISS174
count
Reference frequency

(b)

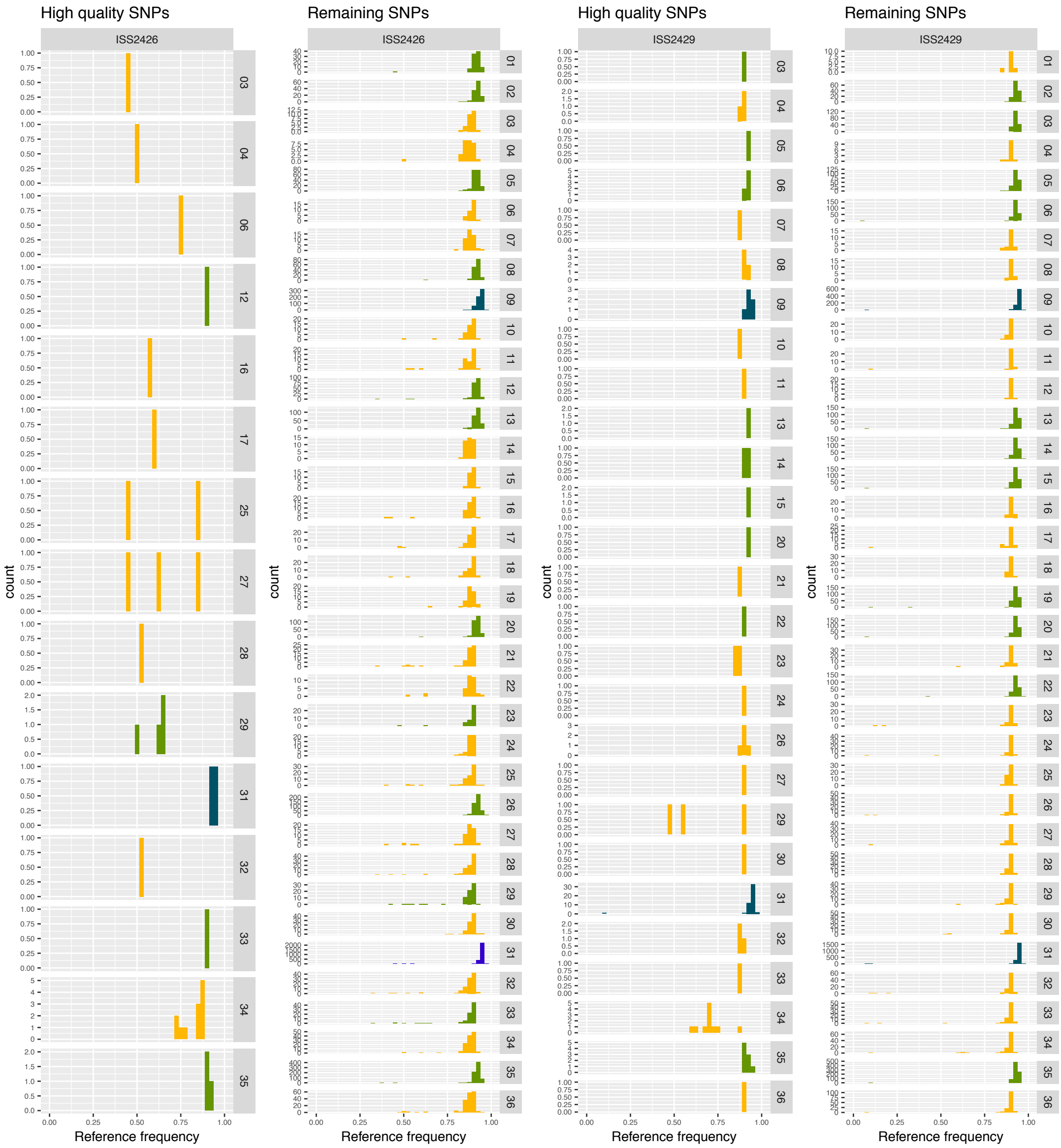
High quality SNPs
ISS2426
Remaining SNPs
ISS2426
High quality SNPs
ISS2429
Remaining SNPs
ISS2429
count
Reference frequency

(c)

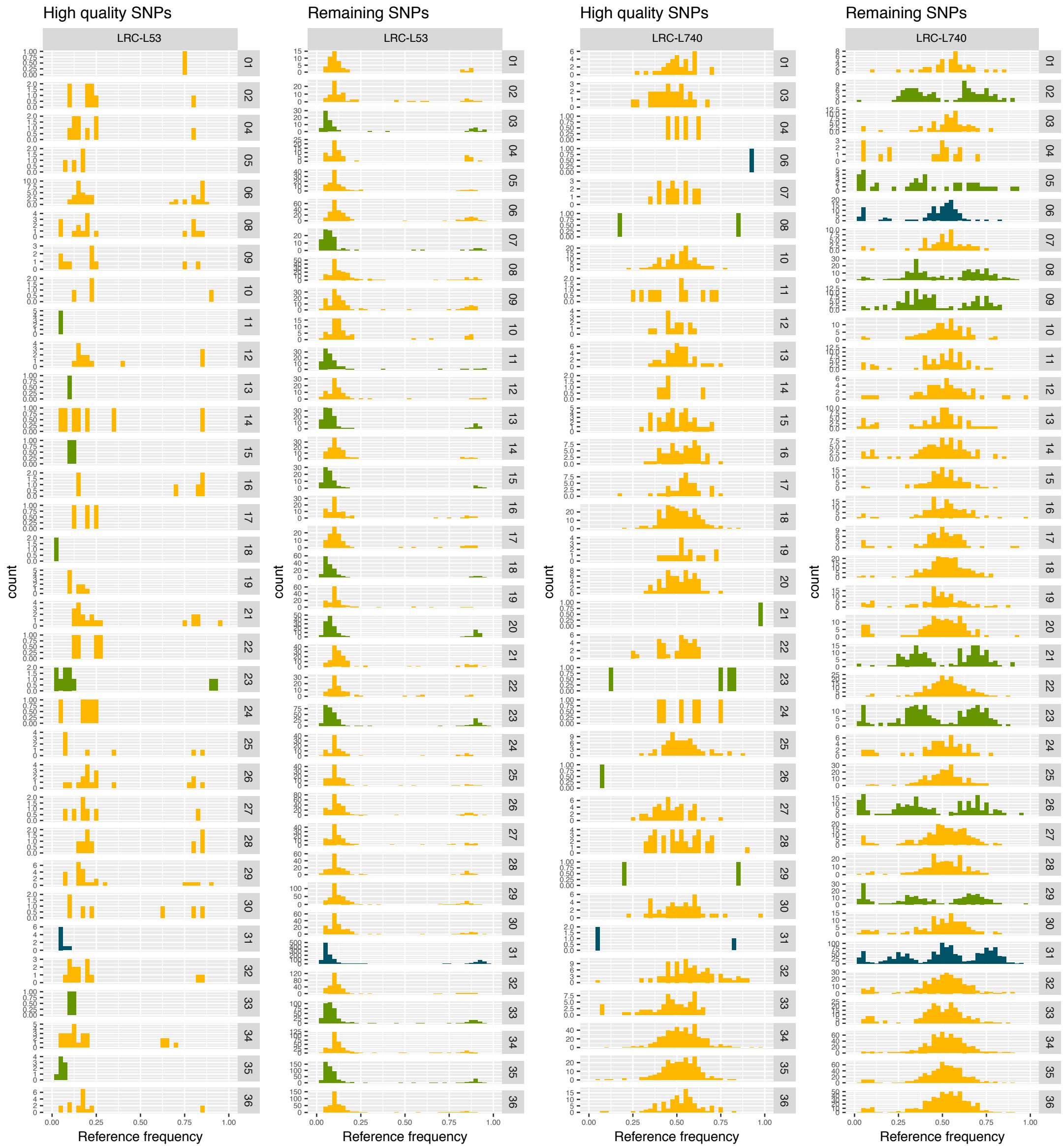
High quality SNPs
LRC-L53
Remaining SNPs
LRC-L53
High quality SNPs
LRC-L740
Remaining SNPs
LRC-L740
count
Reference frequency
0.00 0.25 0.50 0.75 1.00

(d)

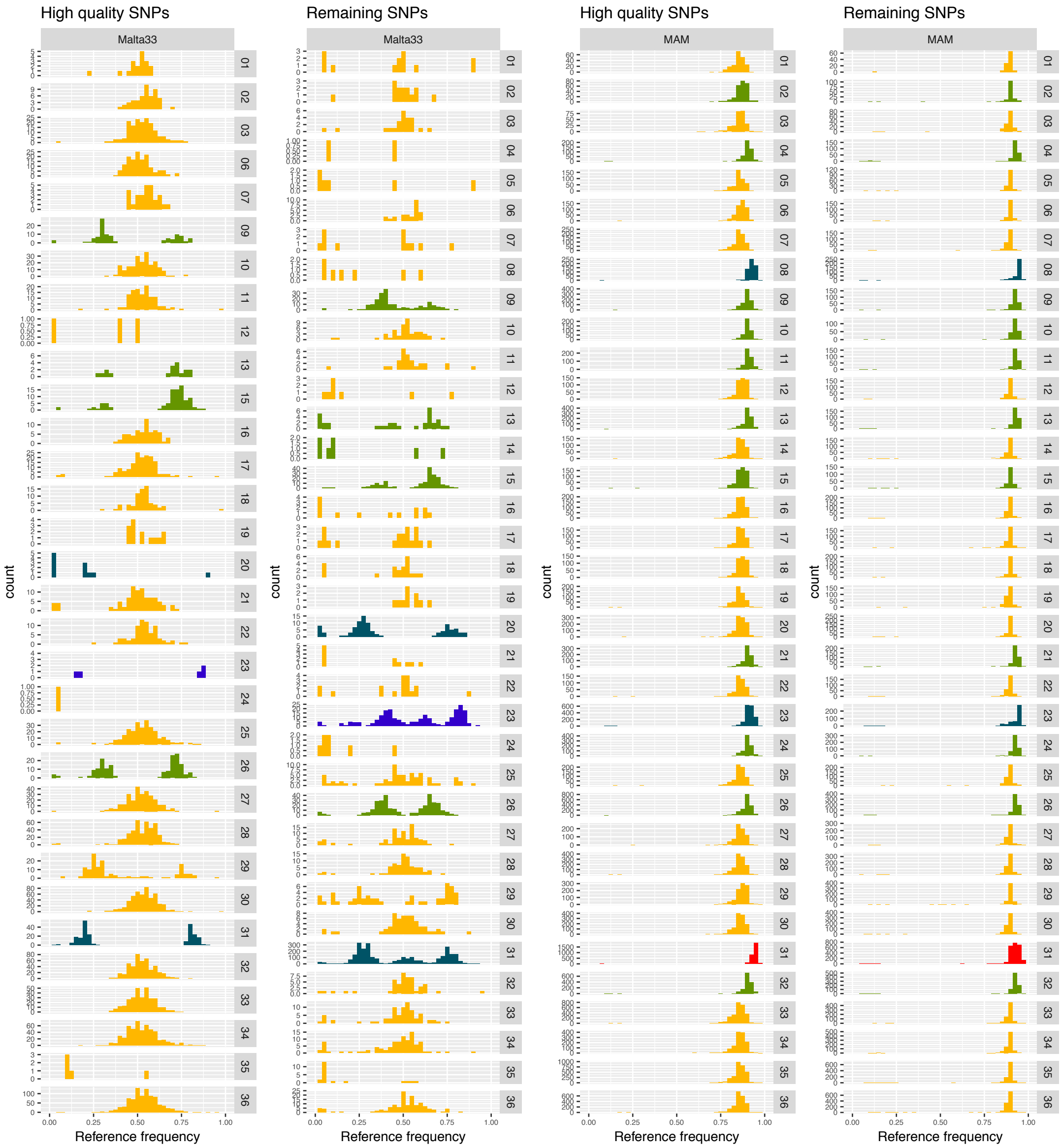

High quality SNPs
Remaining SNPs
High quality SNPs
Remaining SNPs
Malta33
Malta33
MAM
MAM
count
Reference frequency

(e)

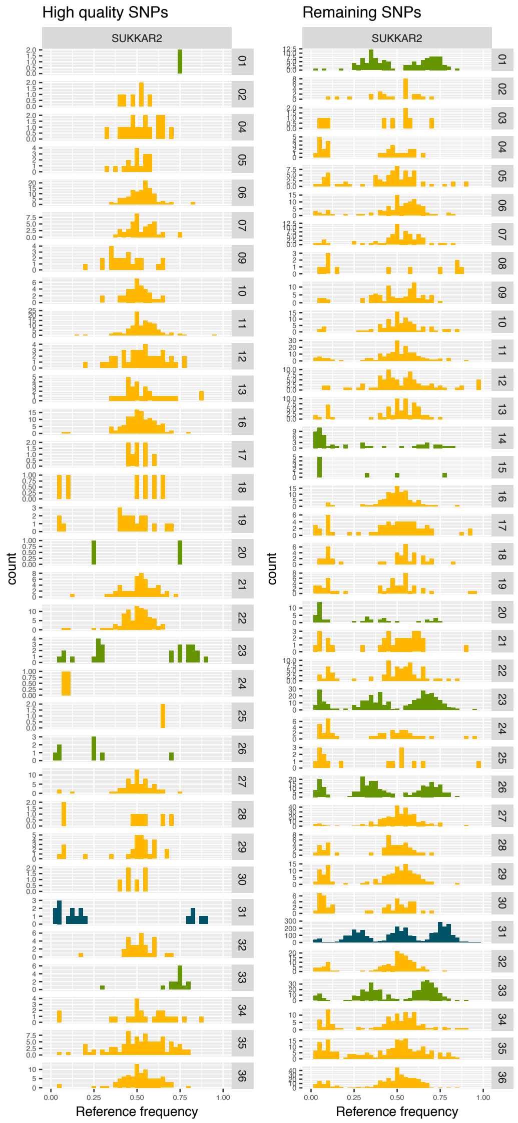

(f)

Figure 13: Verification of skewed allele frequency spectra in a subset of isolated strains. For all 11 samples, where sample allele frequency distributions indicated mixed infections heterozygous SNPs were filtered for highest quality SNPs (SNP calling quality of 99 and presence of the alternate allele in at least five other isolate strains as homozygous call). For all samples highest and remaining heterozygous SNP frequency distributions within the respective sample are shown. Signals for the suggested highly homozygous sub-clones BPK157A1, Inf152, ISS174, ISS2426, ISS2429, LRC-L53 and MAM, and the sub-clones BUMM3, LRC-L740, Malta33 and SUKKAR2 with the high-frequency one being heterozygous are mainly supported by highest quality SNPs.

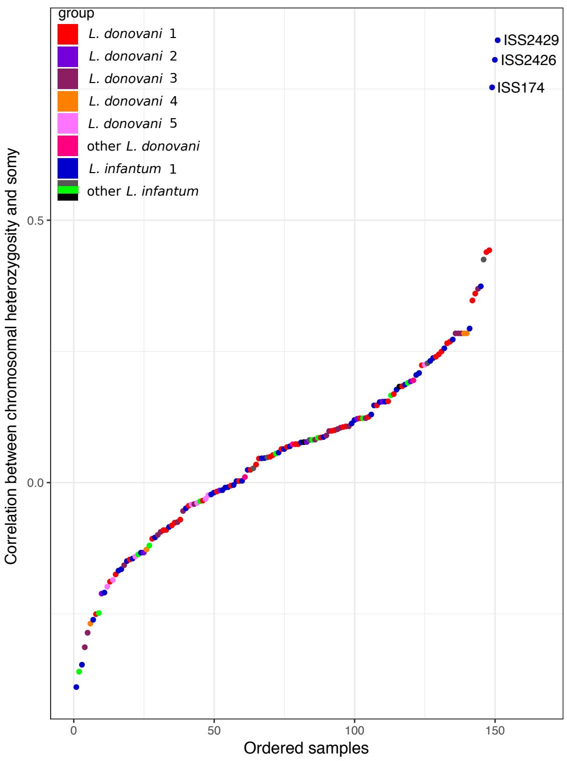

Figure 14: Correlation between somies and heterozygosities across chromosomes. For all 151 isolates aneuploidy profiles were correlated with respective chromosome-specific heterozgosity values. Names of isolates with significant correlations (Spearman, FDR <= 0.001) are printed next to the respective correlation value.

#### 1.6 Population genomic characterisation of the groups

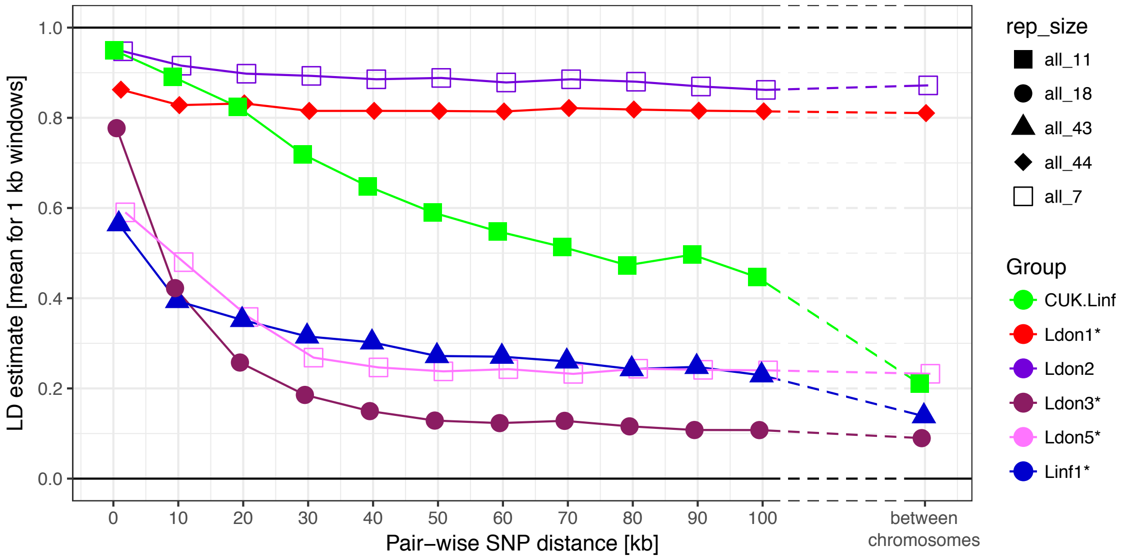

Figure 15: LD decay with genomic distance. LD decay was measured for the six largest (sub-)groups removing isolates that were identified as putative strain mixtures (indicated by *; see Material & Methods). Shown are means of 1 kb wide windows for SNP pairs of the stated genomic distance. For LD estimates between chromosomes, 100 SNPs were randomly sampled per chromosome and means across all pair-wise combinations between chromosome are shown. This procedure was done twice independently but as differences between both such replicates were negligible only the results of one replicate are shown. Samples sizes used for LD estimates vary between groups depending on the group size.

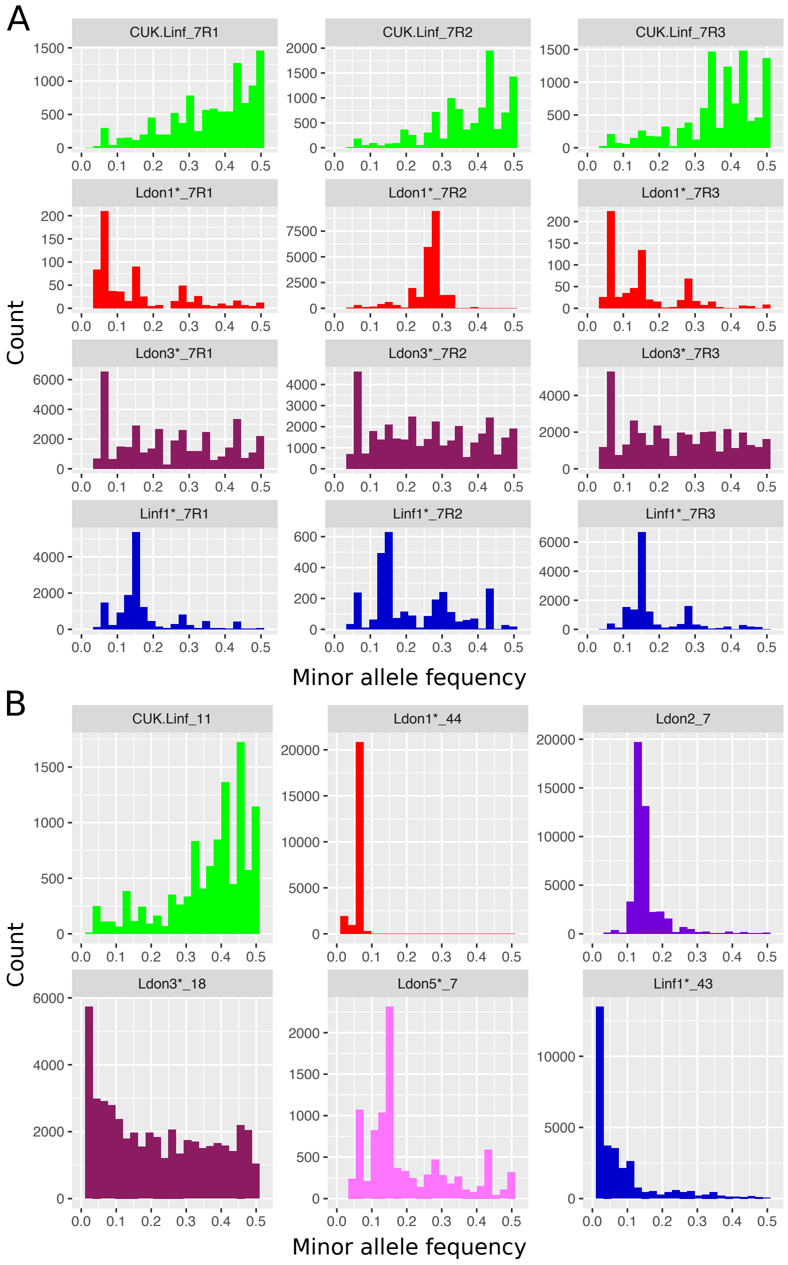
A
CUK.Linf_7R1
CUK.Linf_7R2
CUK.Linf_7R3
Ldon1*_7R1
Ldon1*_7R2
Ldon1*_7R3
Ldon3*_7R1
Ldon3*_7R2
Ldon3*_7R3
Linf1*_7R1
Linf1*_7R2
Linf1*_7R3
Count
Minor allele fequency
B
CUK.Linf_11
Ldon1*_44
Ldon2_7
Ldon3*_18
Ldon5*_7
Linf1*_43
Count
Minor allele fequency

Figure 16

Figure 16: Folded site frequency spectra of the six largest groups. A) The six largest (sub-)groups were randomly sub-sampled to seven samples in three pseudo-replicates and SFS are shown. B) SFS are shown for the six largest (sub-)groups including all available samples. Group names are indicated at the top of each plot. Asterisks indicate that samples that were identified as clone mixtures were removed (see Material & Methods). Sample size and replicate number if applicable are written next to the group names.

#### 1.7 Fixation index and marker SNPs for the different groups

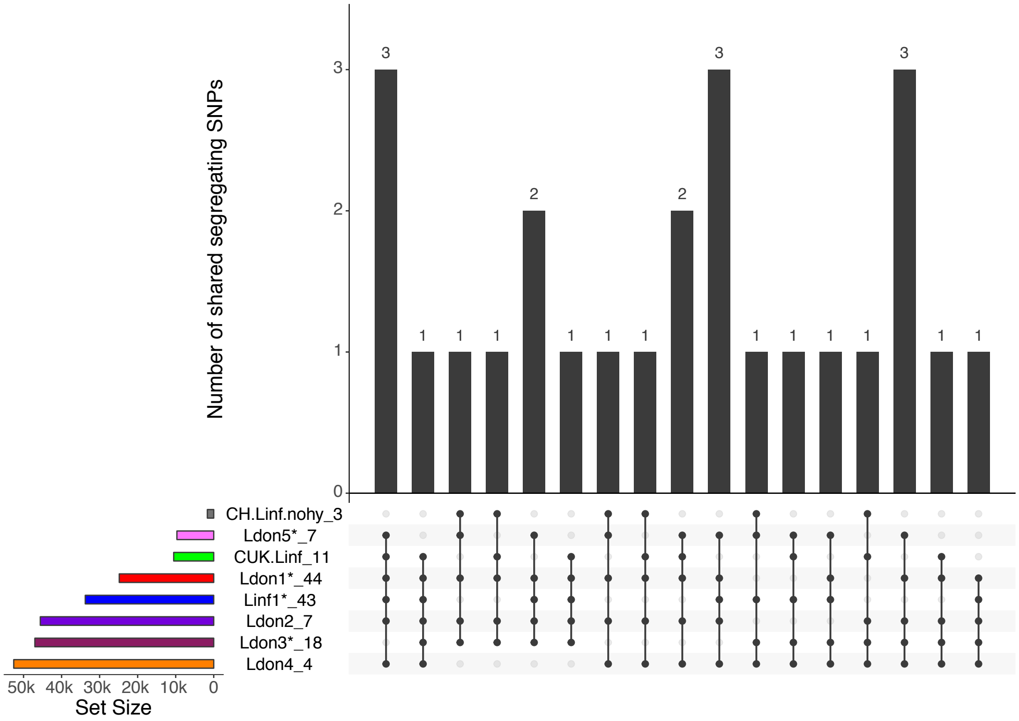

Figure 17: Polymorphism sharing between (sub-)groups. The histogram lists the number of SNPs that are segregating in multiple groups indicated by a black dot in the panel below. Polymorphism sharing between groups is only shown for sites that are shared by at least five of the eight groups. The histogram on the left shows the number of segregating polymorphisms in each group individually.

#### 1.8 Copy number variation

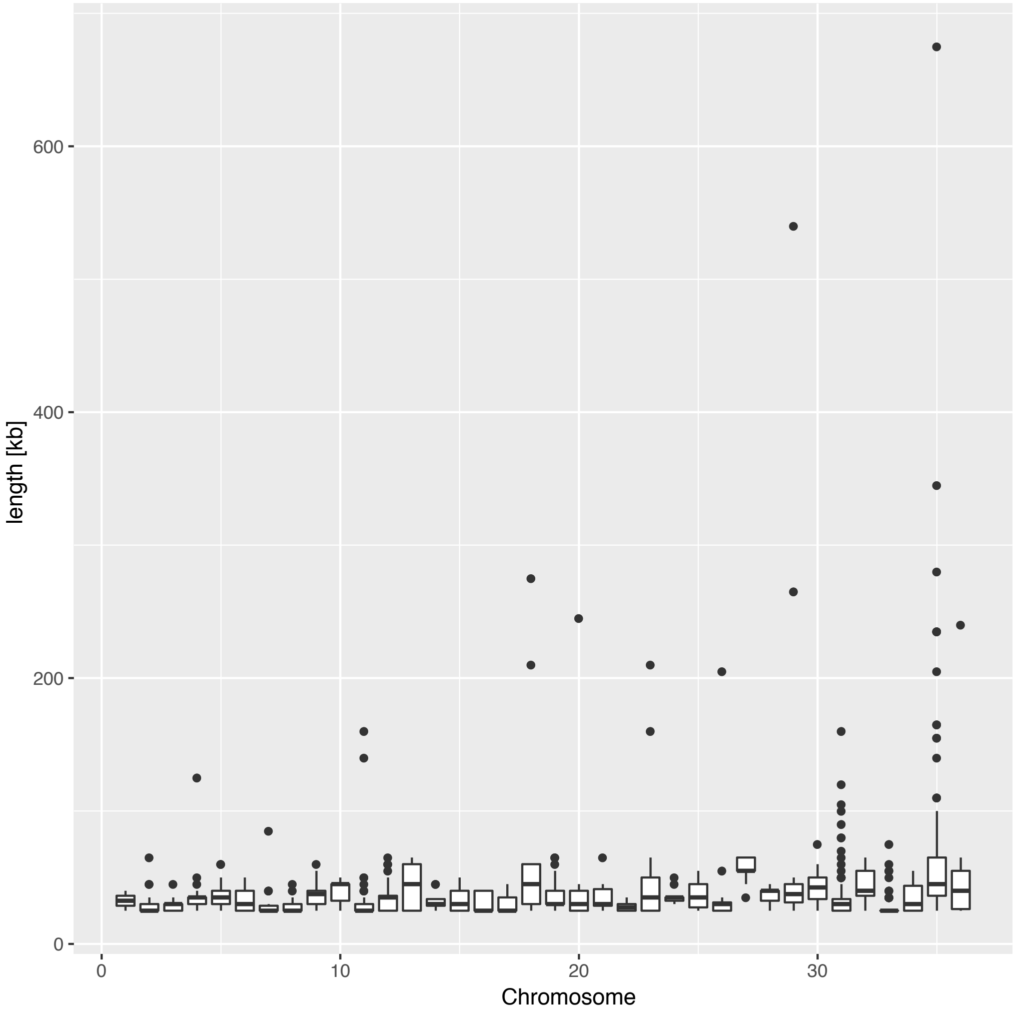

Figure 18: Length distribution of large CNVs by chromosome. Large CNVs were called using a minimum length threshold of 25 kb (see Material & Methods).

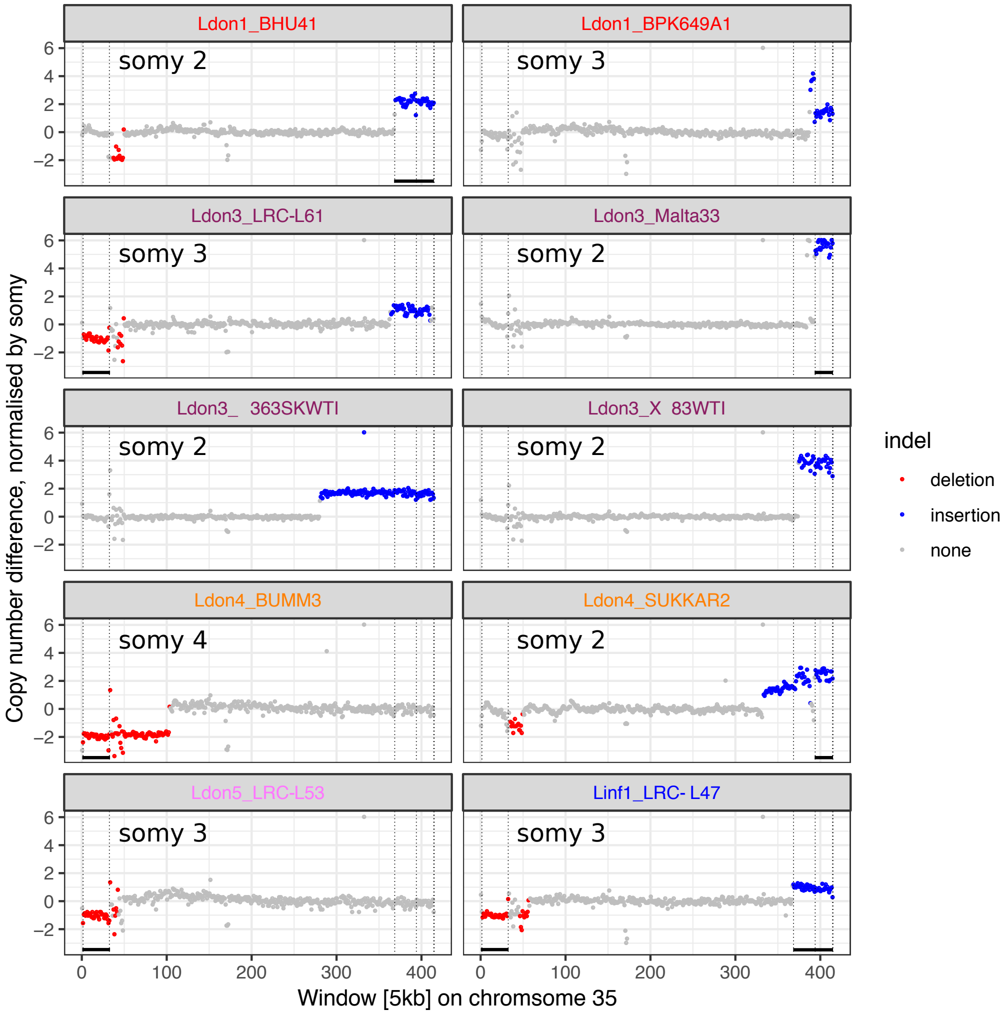

Figure 19: Most chromosome scale CNVs are located on chromosome 35. Shown are all samples of chromosome 35 that harbour at least one chromosome-scale CNV (>100 kb). Genome coverage for 5 kb windows was normalised by the overall chromosome somy and coloured in red and blue for duplications and deletions, respectively. The overall somy by sample for chromosome 35 is indicated in each plot. Vertical lines mark indel boundaries and horizontal black bars below indicate indels with shared identical boundaries between samples. Group origin of the different samples is indicated by the group colours used throughout this study.

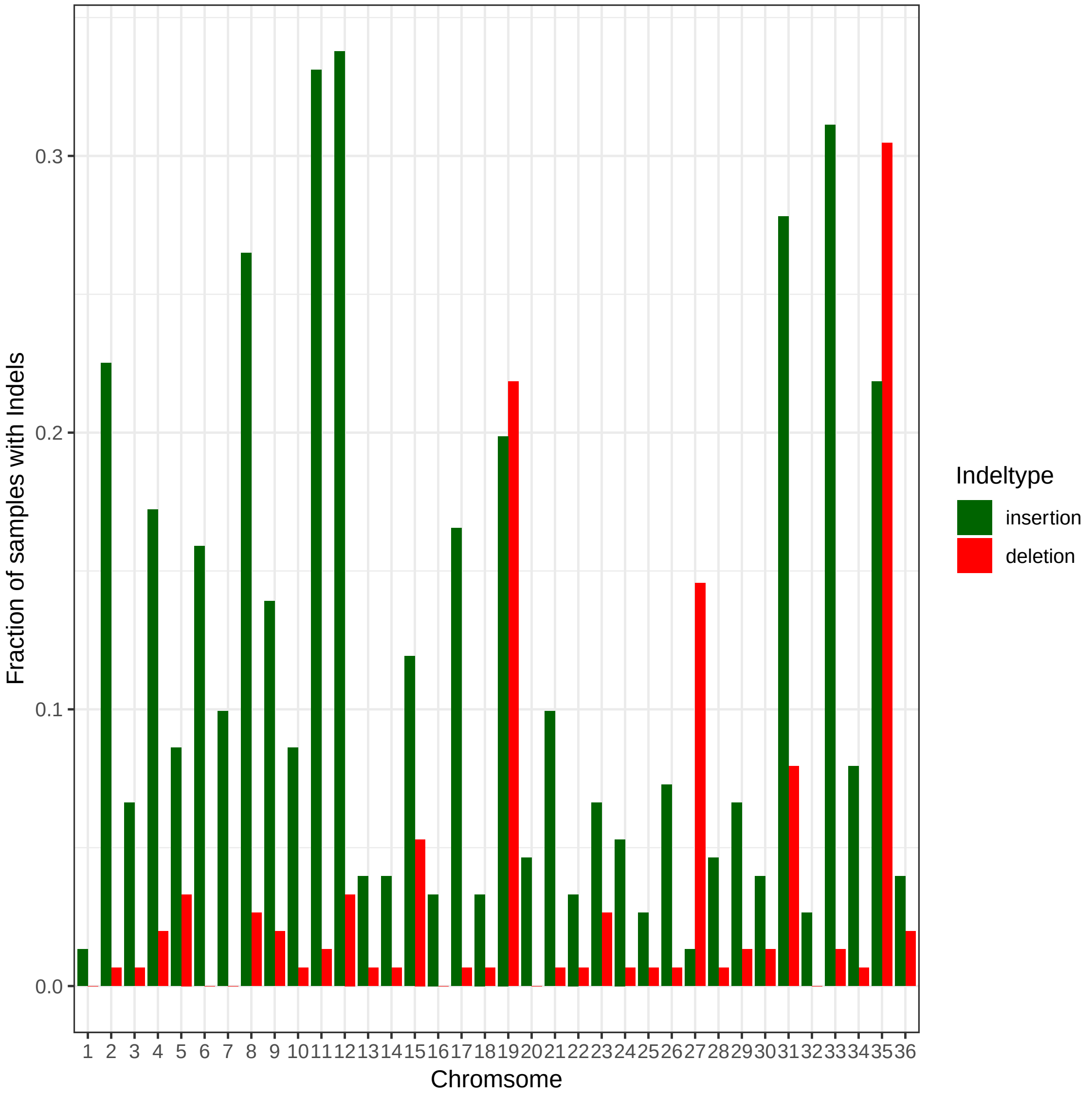

Figure 20: Fraction of large CNVs across chromosomes. Shown is the fraction of all 151 samples that contain at least one large copy number variant (>= 25 kb; see Material & Methods) of the respective type for each chromosome.

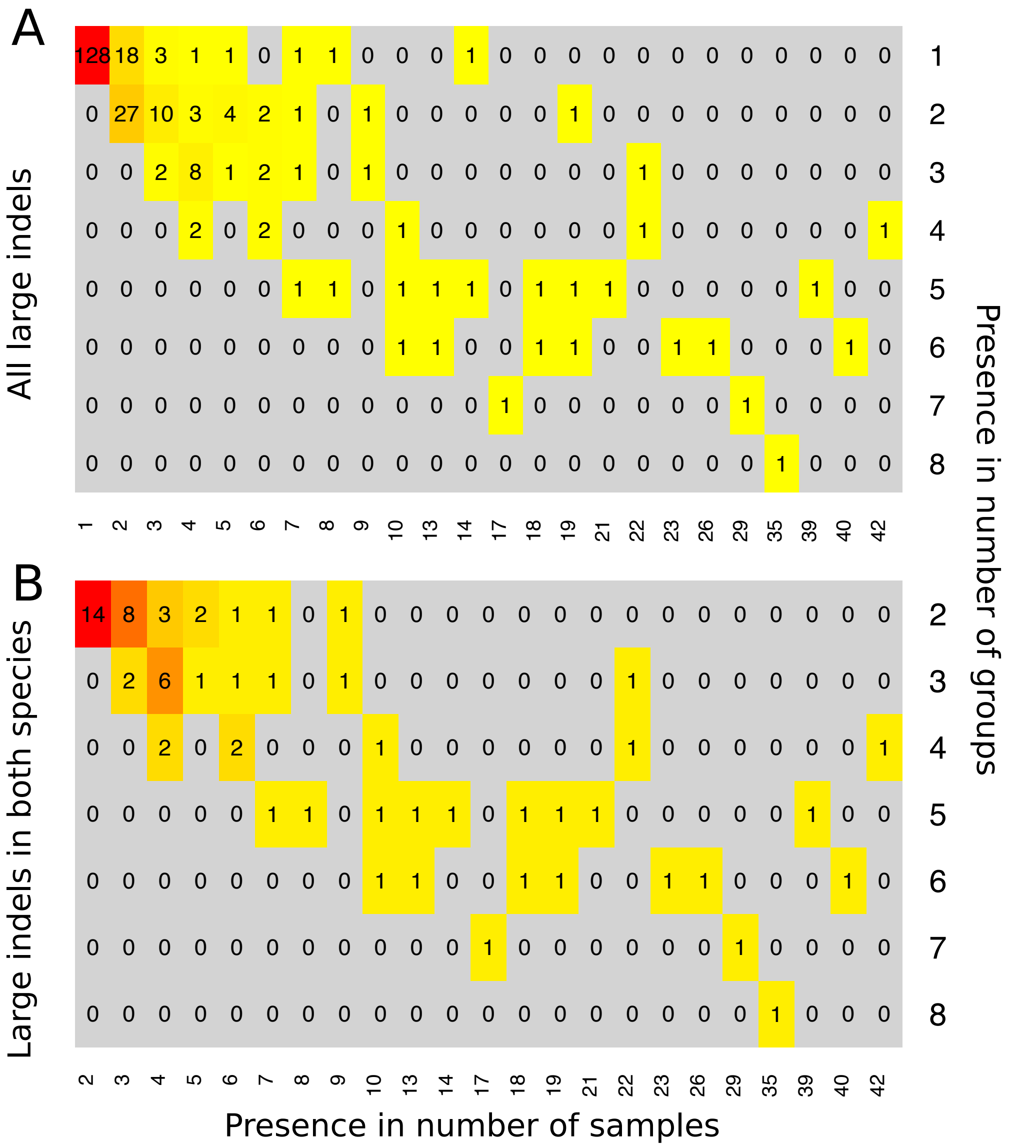

Figure 21: Large CNVs shared across samples and groups. Sharing of large CNVs (>=25 kb) is shown between samples and groups. A) All large CNVs identified across all 151 isolates. B) All large CNVs that have been found in both species, *L. donovani* and *L. infantum*.

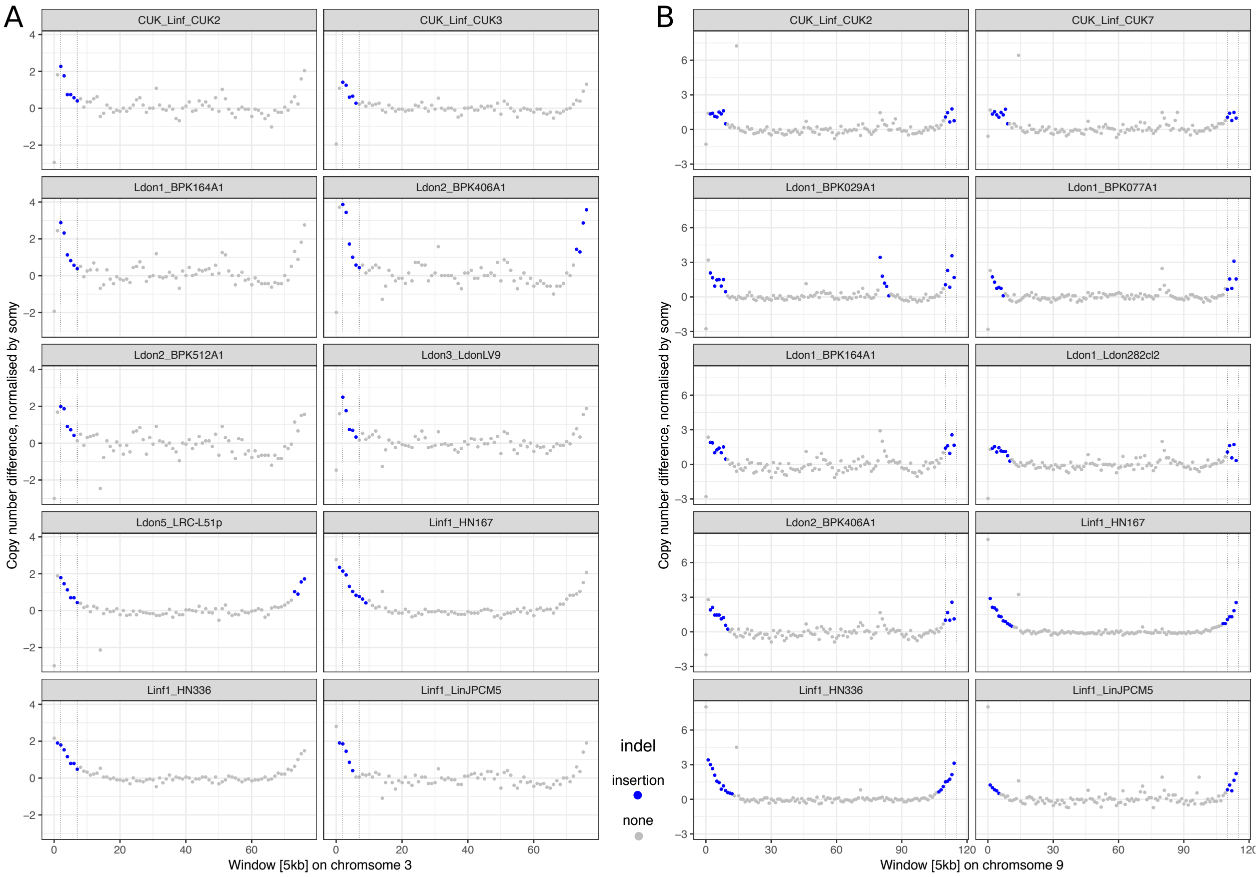

Figure 22: Increased coverage of samples towards chromosome ends. Samples are shown with called duplications at chromosome ends that show a gradual coverage increase. Plots show median window coverage across 5 kb windows. Called duplications are indicated by blue dots and boundaries are indicated by vertical bars. A) Examples for chromosome 3. B) Examples for chromosome 9.

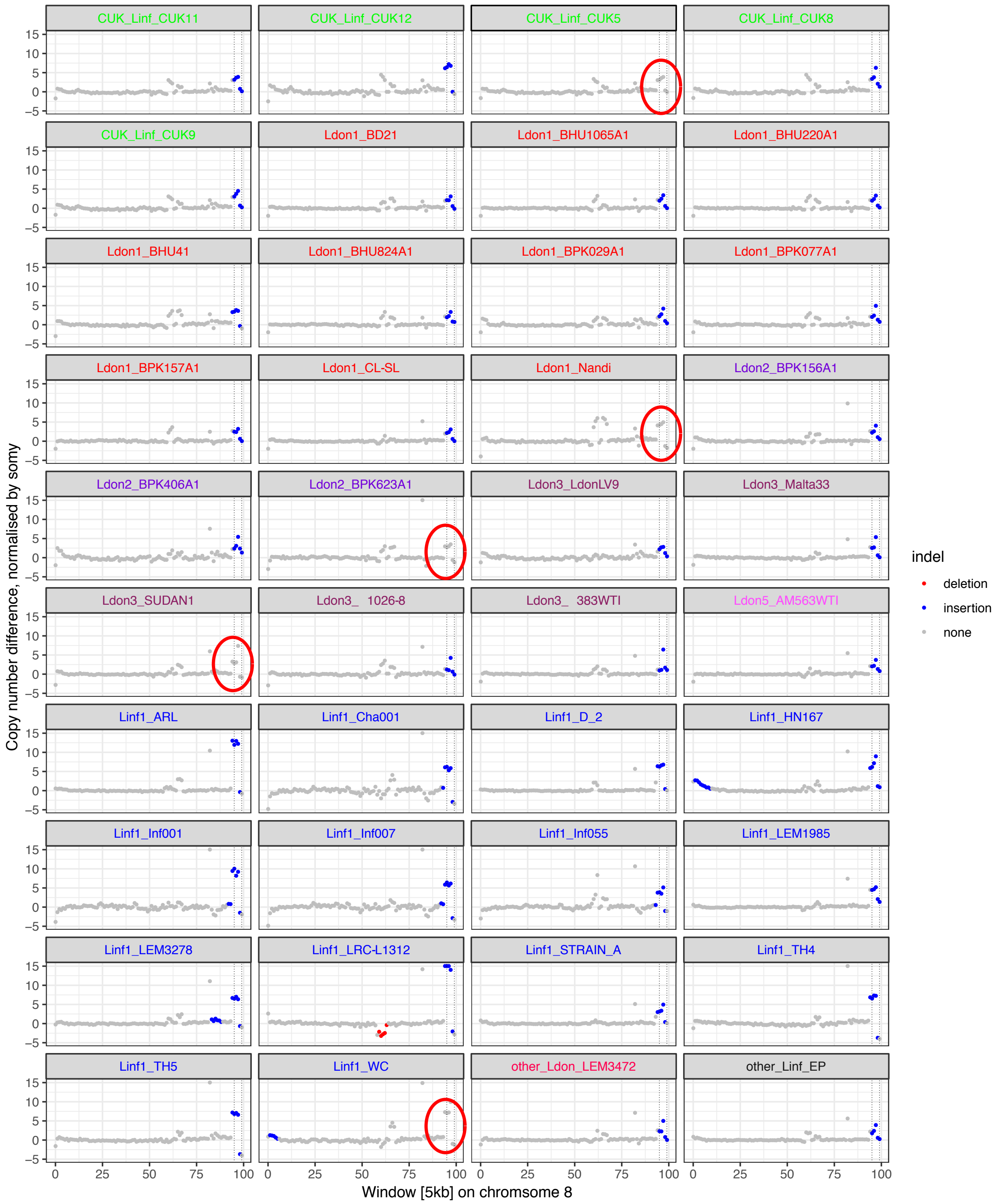
CUK_Linf_CUK11
CUK_Linf_CUK12
CUK_Linf_CUK5
CUK_Linf_CUK8
CUK_Linf_CUK9
Ldon1_BD21
Ldon1_BHU1065A1
Ldon1_BHU220A1
Ldon1_BHU41
Ldon1_BHU824A1
Ldon1_BPK029A1
Ldon1_BPK077A1
Ldon1_BPK157A1
Ldon1_CL-SL
Ldon1_Nandi
Ldon2_BPK156A1
Ldon2_BPK406A1
Ldon2_BPK623A1
Ldon3_LdonLV9
Ldon3_Malta33
Ldon3_SUDAN1
Ldon3_ 1026-8
Ldon3_ 383WTI
Ldon5_AM563WTI
Linf1_ARL
Linf1_Cha001
Linf1_D_2
Linf1_HN167
Linf1_Inf001
Linf1_Inf007
Linf1_Inf055
Linf1_LEM1985
Linf1_LEM3278
Linf1_LRC-L1312
Linf1_STRAIN_A
Linf1_TH4
Linf1_TH5
Linf1_WC
other_Ldon_LEM3472
other_Linf_EP
Copy number difference, normalised by somy
Window [5kb] on chromsome 8
indel
deletion
insertion
none

Figure 23

Figure 23: Indication of a putative assembly error in the reference genome. A common duplication of 25 kb on chromosome 8, position $470 - 495$ kb, was found in 35 samples across 8 different groups (indicated by vertical bars and highlighted in blue). When inspecting remaining samples, however, a copy number increase was also present in all other 116 remaining samples, which failed to meet the CNV calling threshold. Five of these 116 samples are also shown and the non-called copy number increase is indicated by a red circle. As the copy number increase varies between samples, this regions may still be copy number variable between isolates.

#### 1.9 Genetic variation for known drug resistance loci

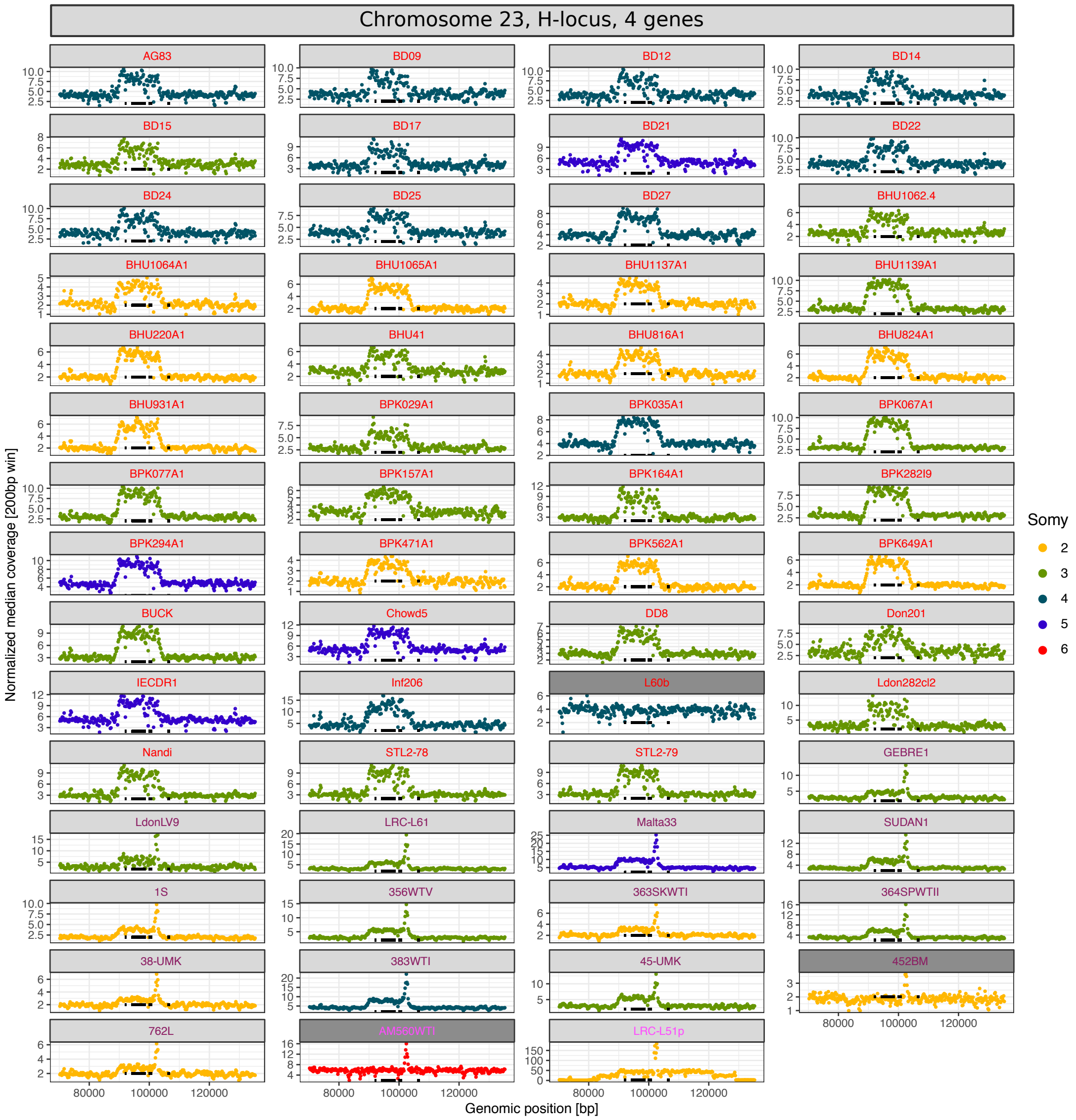

(a)

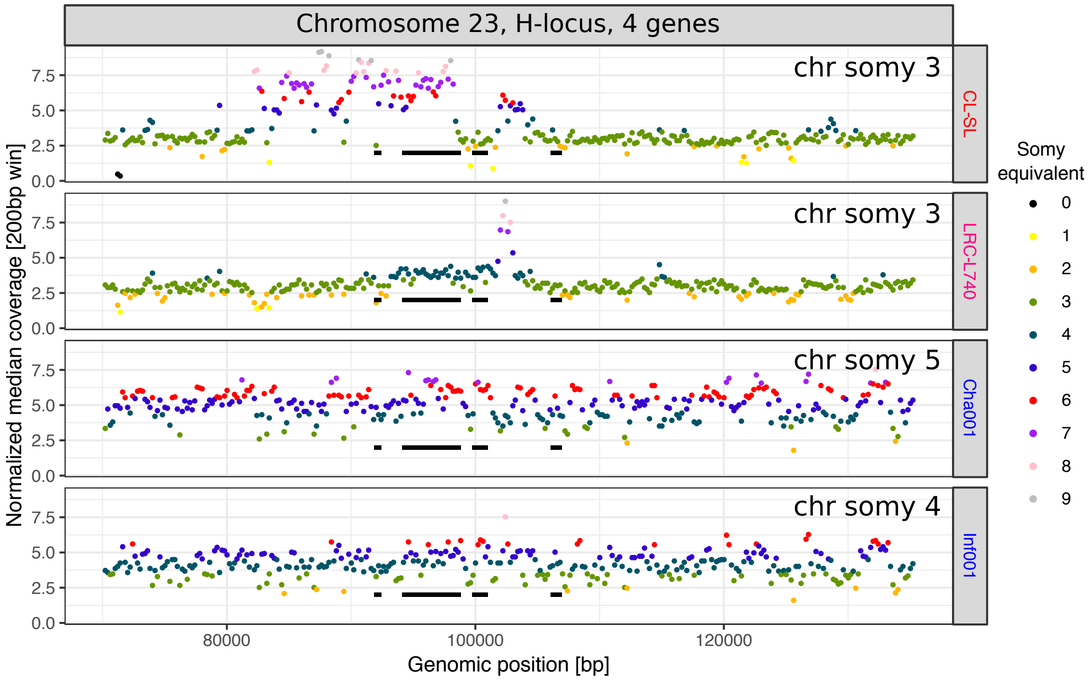

(b)

Figure 24: Copy number increase at the H-locus. Shown are coverage plots across isolates for the H-locus on chromosome 23 highlighting the four genes (YIP1, MRPA, argininosuccinate synthase, PTR1) present at this locus by black, vertical bars. Isolate names are coloured by the groups colours used throughout this study. a) Copy number variation is shown for all 37% of isolates that have a copy number increase of at least 3 genes at the H-locus. Coverages of each isolate are coloured by the somy of chromosome 23 in the respective isolate. For comparison for each group, the coverage of one isolate with no copy number increase is also plotted (dark grey headers). b) Copy number variation of all isolates that show a copy number increase in only two of the associated genes. Window-specific coverages are coloured by the somy equivalent of the respective coverage. Somies of chromosome 23 for each isolates are indicated in the respective row.

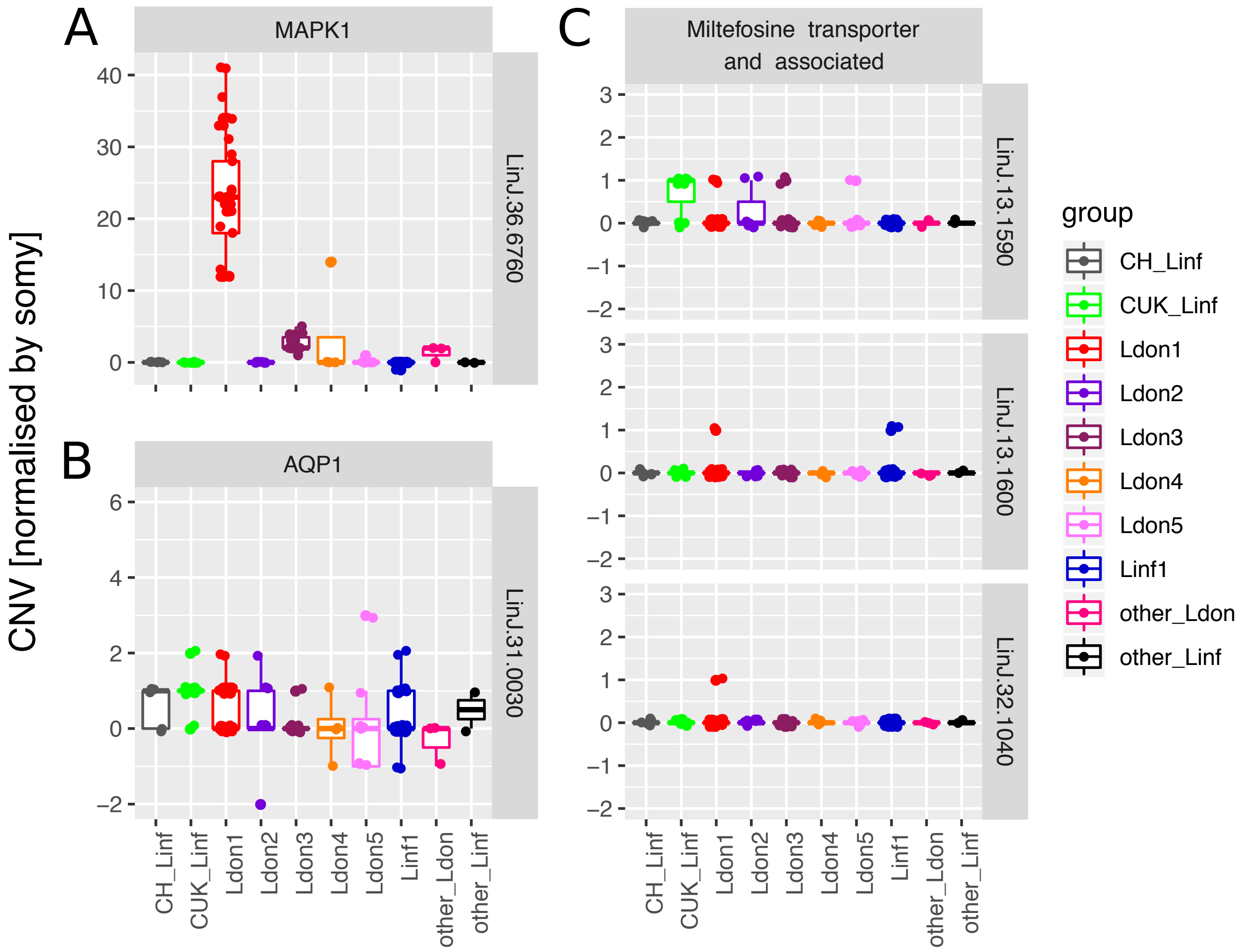

Figure 25: Copy number variation of putative drug resistance genes. Gene copy number for all four genes and all 151 samples is shown across 10 different (sub-)groups for three different loci putatively involved in drug resistance. A) MAPK1 (LinJ.36.6760) and B) AQP1 (LinJ.31.0030) genes are putatively involved antimony drug resistance. C) The Miltefosine transporter (LinJ.13.1590), the adjacent gene (LinJ.13.1600) and the Ros3 (LinJ.32.1040) gene are putatively involved in Miltefosine resistance.

#### 1.10 Population and species specific selection

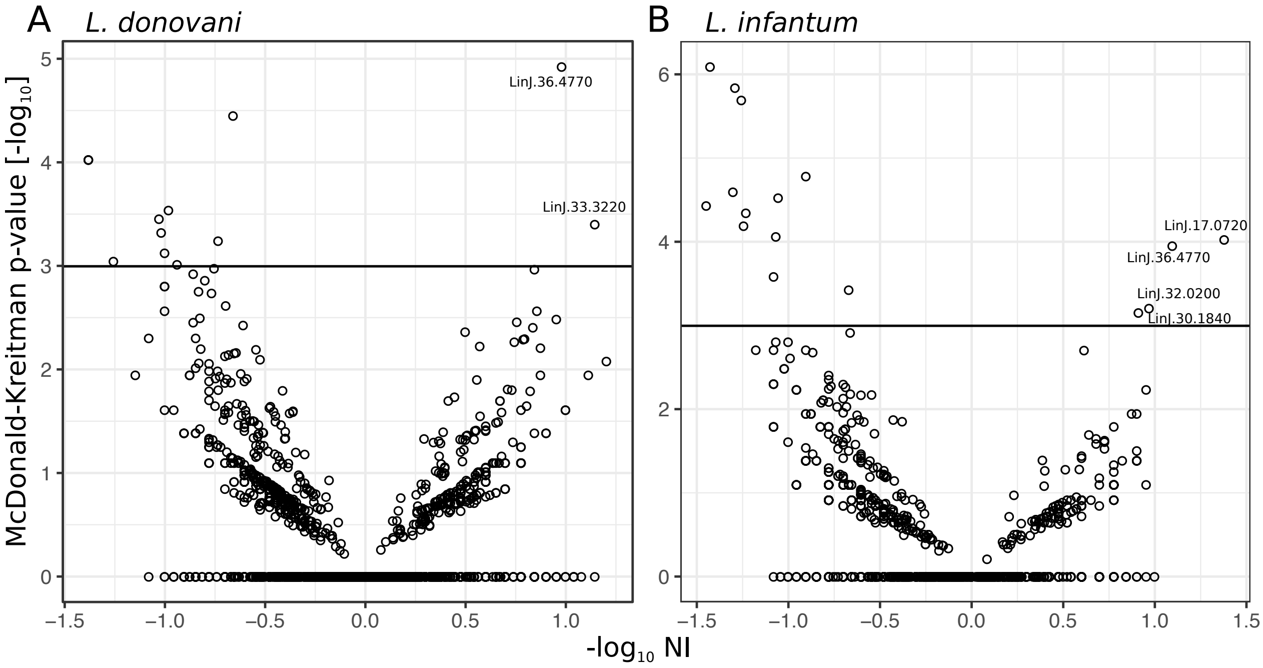

Figure 26: Measures of adaptive evolution. Species species-specific evolution is measured for A) *L. donovani* and B) *L. infantum*. Each point represents the neutrality index (NI) and the associated p-value of the McDonald-Kreitman test for each of 8,234 genes. The horizontal line represents the $-log_{10}$ value equivalent to a p-value of 0.05. None of the shown values passes multiple testing correction.

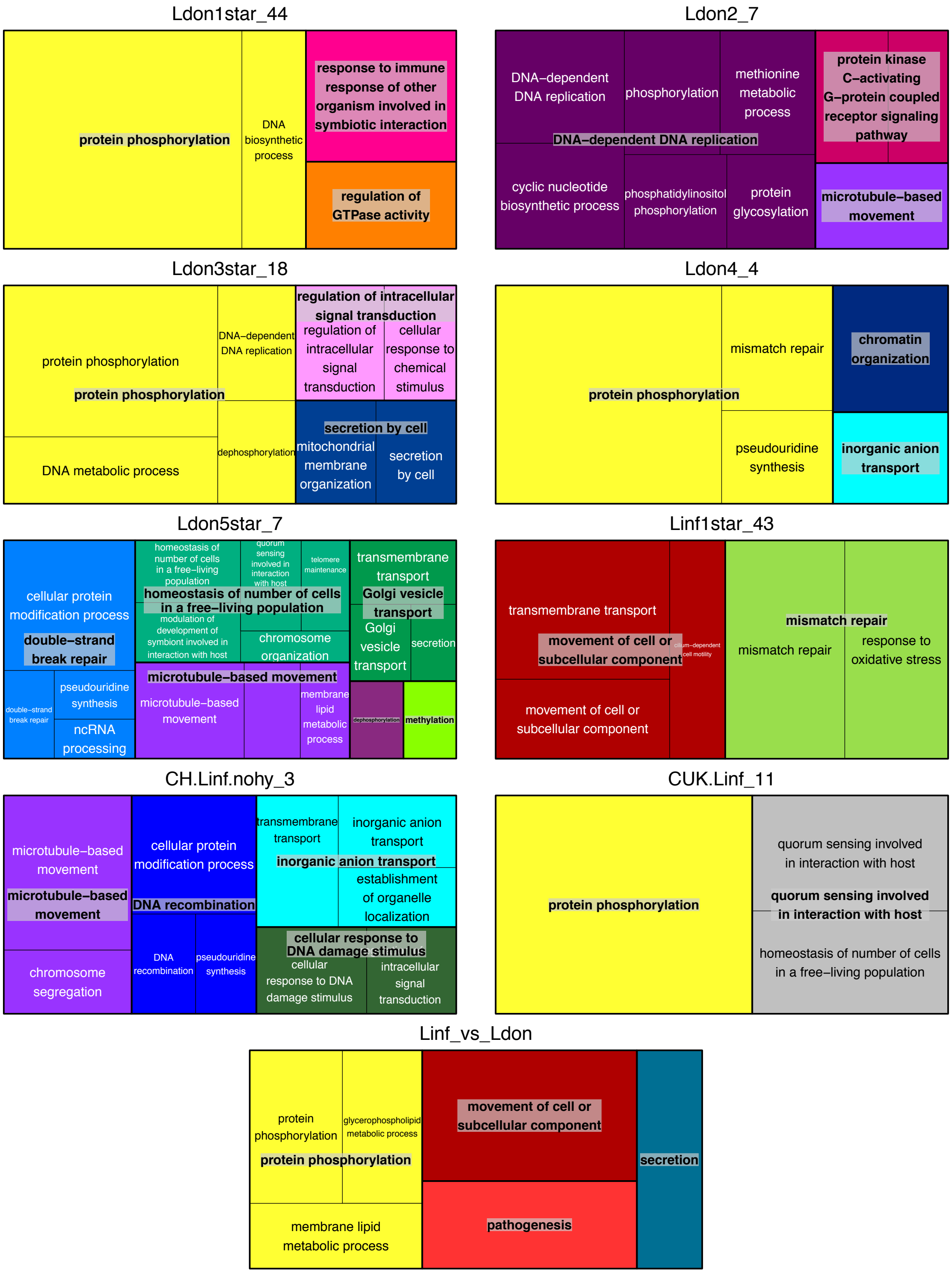
Ldon1star_44
protein phosphorylation
DNA biosynthetic process
response to immune response of other organism involved in symbiotic interaction
regulation of GTPase activity
Ldon2_7
DNA−dependent DNA replication
phosphorylation
methionine metabolic process
DNA−dependent DNA replication
cyclic nucleotide biosynthetic process
phosphatidylinositol phosphorylation
protein glycosylation
protein kinase C−activating G−protein coupled receptor signaling pathway
microtubule−based movement
Ldon3star_18
protein phosphorylation
protein phosphorylation
DNA−dependent DNA replication
DNA metabolic process
dephosphorylation
regulation of intracellular signal transduction
regulation of intracellular signal transduction
cellular response to chemical stimulus
secretion by cell
mitochondrial membrane organization
secretion by cell
Ldon4_4
protein phosphorylation
mismatch repair
pseudouridine synthesis
chromatin organization
inorganic anion transport
Ldon5star_7
cellular protein modification process
double−strand break repair
double−strand break repair
pseudouridine synthesis
ncRNA processing
homeostasis of number of cells in a free−living population
quorum sensing involved in interaction with host
telomere maintenance
homeostasis of number of cells in a free−living population
modulation of development of symbiont involved in interaction with host
chromosome organization
transmembrane transport
Golgi vesicle transport
Golgi vesicle transport
secretion
microtubule−based movement
microtubule−based movement
membrane lipid metabolic process
dephosphorylation
methylation
Linf1star_43
transmembrane transport
movement of cell or subcellular component
cilium−dependent cell motility
movement of cell or subcellular component
mismatch repair
mismatch repair
response to oxidative stress
CH.Linf.nohy_3
microtubule−based movement
microtubule−based movement
chromosome segregation
cellular protein modification process
DNA recombination
DNA recombination
pseudouridine synthesis
transmembrane transport
inorganic anion transport
inorganic anion transport
establishment of organelle localization
cellular response to DNA damage stimulus
cellular response to DNA damage stimulus
intracellular signal transduction
CUK.Linf_11
protein phosphorylation
quorum sensing involved in interaction with host
quorum sensing involved in interaction with host
homeostasis of number of cells in a free−living population
Linf_vs_Ldon
protein phosphorylation
glycerophospholipid metabolic process
protein phosphorylation
membrane lipid metabolic process
movement of cell or subcellular component
pathogenesis
secretion

Figure 27

Figure 27: Gene ontology enrichment of marker genes with putative biological impact. For each group our species GO enrichment, biological process results are shown for genes with at least one moderate or high effect variant according to SNPeff annotation (Tab. S3). Plots show enrichment for the lenient cutoff of p-value$<0.05$ using the the weighted Fisher test statistic (weightFish, topGO, Alexa *et al.*, 2006) using Revigo (Supek *et al.*, 2011). Sizes of rectangles are normalised by absolute log10 p-value. Plot titles indicate the group marker sets or the species comparison, respectively. Stars in the groups names indicate that samples that have previously identified as mixtures of clones have been removed (Tab. 1 B3 & B4). Additionally, hybrids between the major groups (Tab. 1 B2) were removed from this analysis. Group sample sizes are indicated at the end of each name.

#### 1.11 Methods: Maxicircle statistics

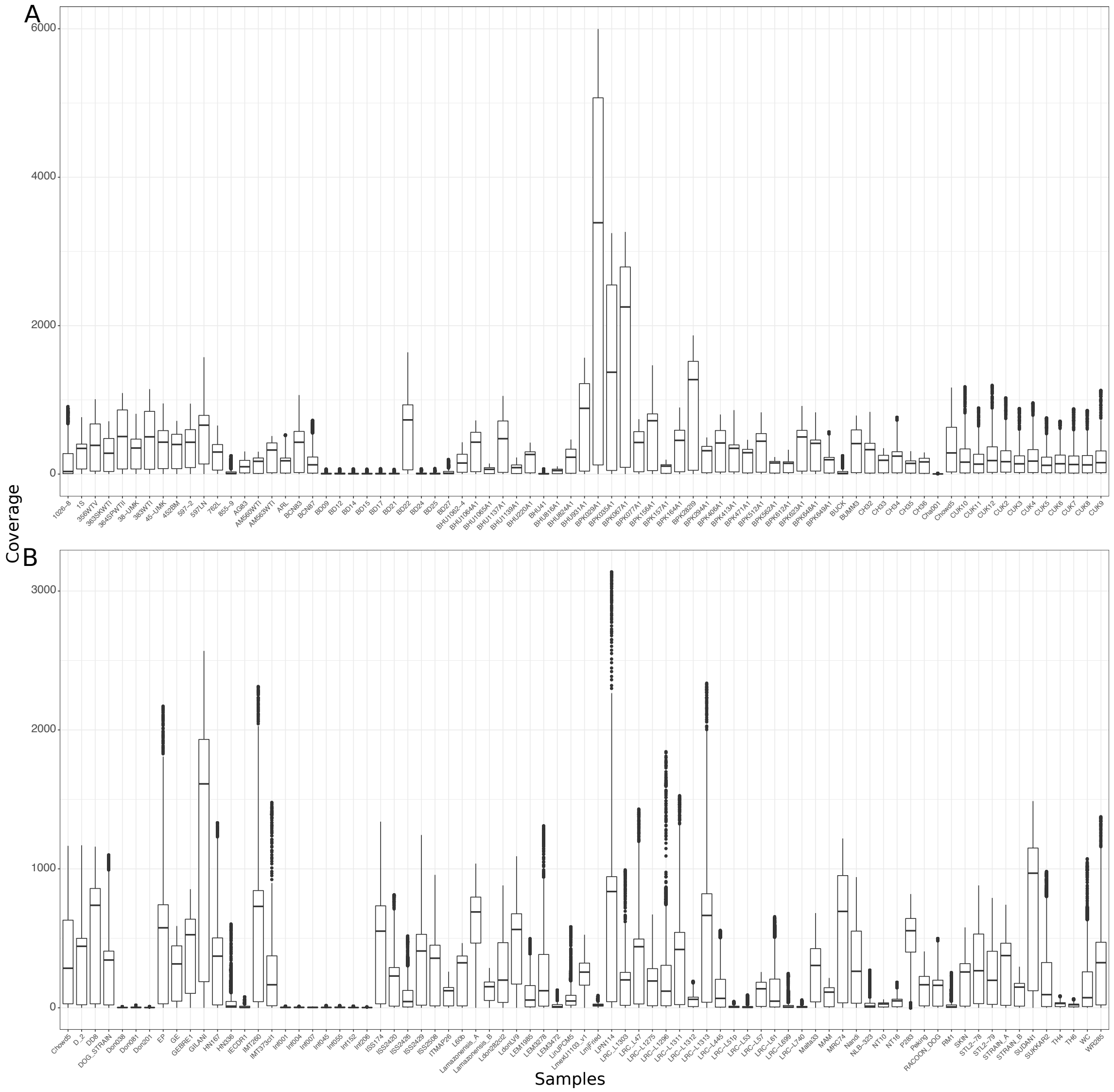

Figure 28: Coverage of maxicircle DNA across isolates. Only isolates 116 of the 151 isolates had a median coverage >= 20 and were used for phylogenetic reconstruction of the maxicircle DNA.

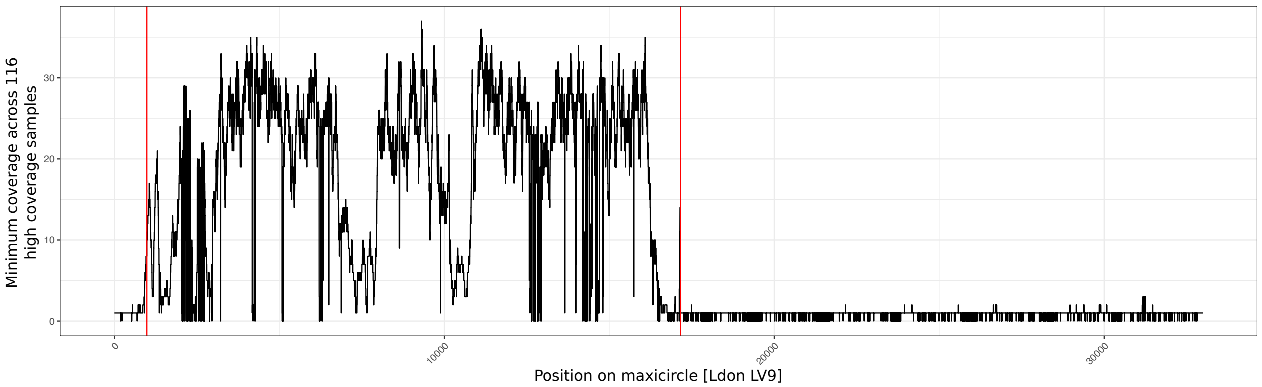

Figure 29: Region of high confidence mapping to the maxicircle DNA across isolates. The minimum coverage across all 116 isolates with a good maxicircle coverage (median coverage >= 20, Fig. 28) is shown along the maxicircle DNA. The region within the red lines was chosen for phylogenetic reconstruction

#### 1.12 Methods: Haplotype-based analysis of hybridisation in CUK isolates

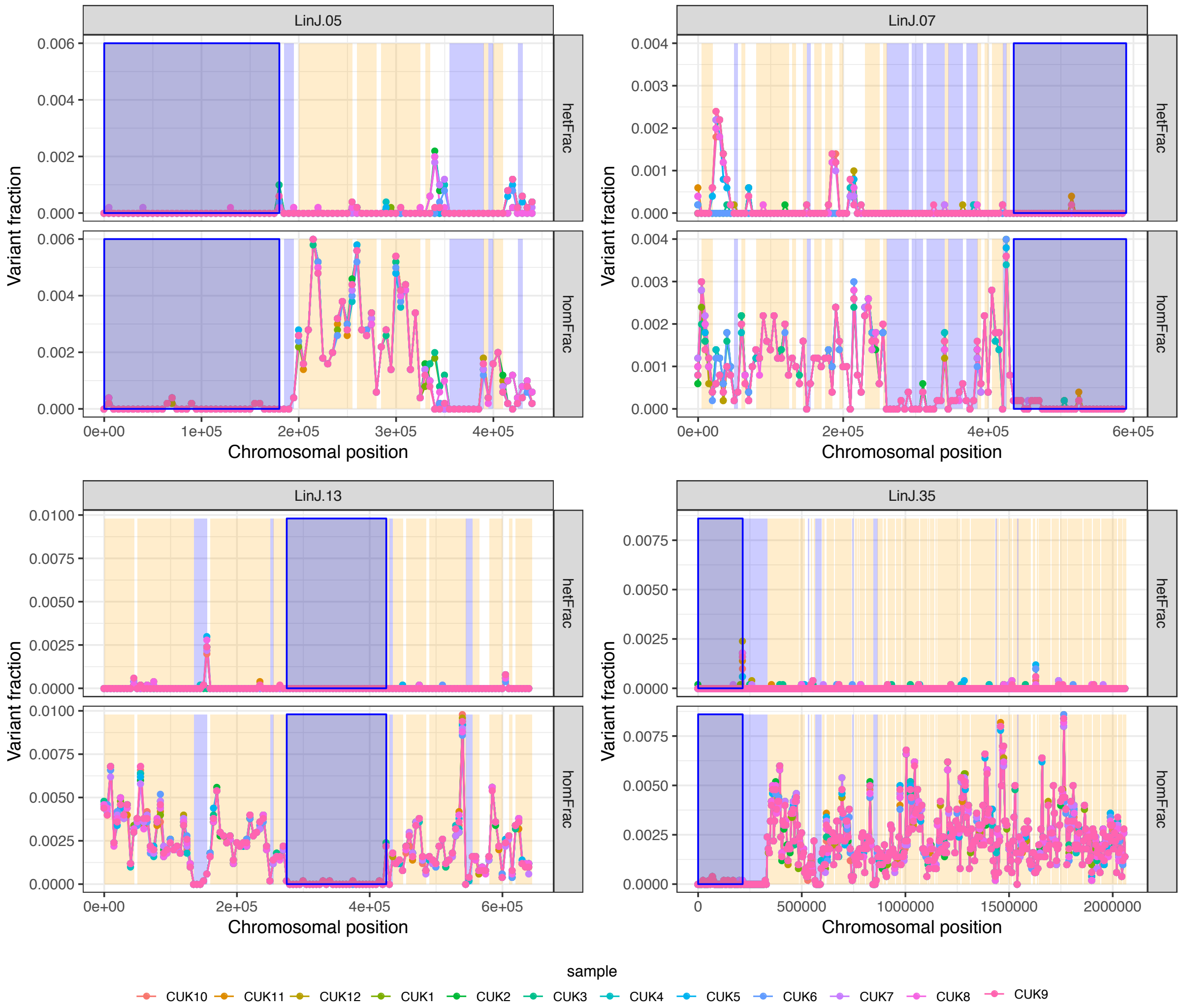

(a)

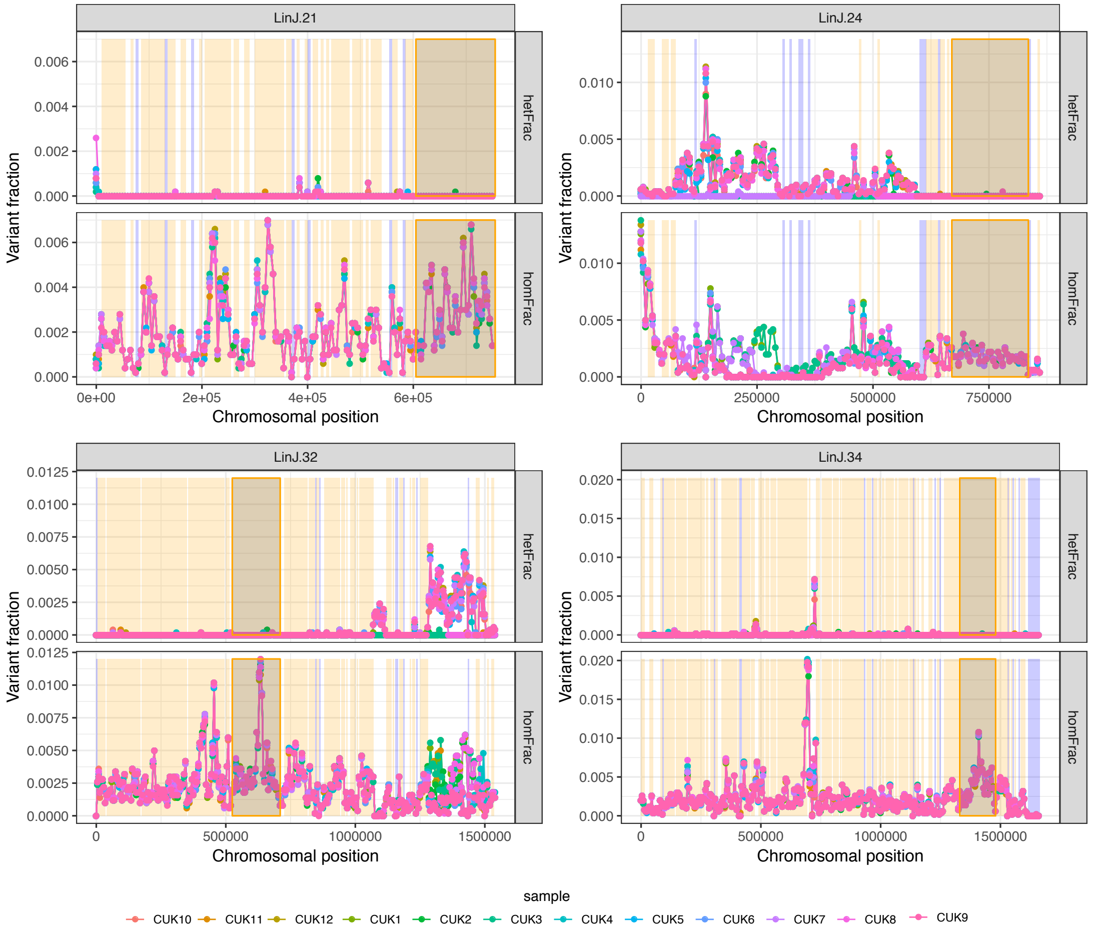

(b)

Figure 30: Genomic regions used for haplotype-based parent identification. Figures show the four largest identified genomic regions specific to either parent used for parent characterisation: (a) JPCM5-like parent and (b) other unknown parent. Each figure shows the fractions of heterozygote SNPs or fixed homozygous SNPs with respect to the JPCM5 reference for each of the 12 isolates from the Cukurova region, Turkey (CUK) (Rogers *et al.*, 2014). Conservatively called regions (see Material & Methods) originating from a specific parent are coloured in blue (JPCM5-like) and orange (other). Genomic regions used for phylogenetic reconstruction are framed.

### 2 Tables

#### 2.1 Table S1

**TableS1.xlsx**
Metadata on all 151 isolates analysed based on whole genome sequencing data.

#### 2.2 Table S2

**TableS2.xlsx**
Samples with identical maxicircle sequences.

#### 2.3 Table S3

**TableS3.xlsx**
Marker SNPs for each individual group and between species group, excluding samples of putatively mixed clone origin, annotated with SNPeff.

#### 2.4 Table S4

**TableS4.xlsx**
SNPs segregating in at least 5 of the 8 identified groups with respective group frequencies and SNPeff annotations.

#### 2.5 Table S5

**TableS5.xlsx**
Metadata on large indels (>= 25 kb) called for each of the 151 samples.

#### 2.6 Table S6

**TableS6.xlsx**
Metadata of the 245 unique (by genomic position) large indels (>= 25 kb) called across all 151 samples.

#### 2.7 Table S7

**TableS7.xlsx**
Meta data on gene copy number across all 8,330 genes and all 151 samples.

#### 2.8 Table S8

**TableS8.xlsx**
Summary of gene copy number analysis including genes with frequent copy number changes and their functional enrichment using GO term enrichment analysis:

- List of all genes with median copy number change across all 151 samples unequal to 0.
- Enriched GO terms across genes that show a median copy number decrease (<=1).
- Enriched GO terms across genes that show a median copy number increase (>=1).
- Enriched GO terms across genes that show a median copy number increase (>=4).
- Listed are genes that contribute to the GO enrichment of genes with a median copy number change <=1, >=1 and >=4 across all samples, respectively.

#### 2.9 Table S9

**TableS9.xlsx**
GO term enrichment for genes including marker SNPs for each of the 8 identified groups and between *L. infantum* and *L. donovani* samples that show predicted effects of moderate to high effect (SNPeff, Cingolani et al., 2012).

#### 2.10 Table S10

**TableS10.xlsx**
Metadata on genes in *L. donovani* and *L. infantum* with a McDonald-Kreitman test p-value <0.05.

### 3 Supplemental Material & Methods

#### 3.1 Somy evaluation based on allele frequency profiles

For isolates with high genome-wide heterozygosity (>=0.004) peaks of allele frequency distributions were estimated for chromosomes with at least 100 SNPs using the the density function (stats package, R programming language). After peak estimation by isolate and chromosome unreasonable peaks were removed, i.e. the ones that are too low (smaller than 0.2 of the highest peak). The estimated peak vector for each chromosome and isolate were then compared to peak distributions expected for the respective chromosome and and isolate somy, e.g. for a diploid, triploid and tetraploid chromosome we expect peaks only at the frequencies 1/2, 1/3 & 2/3 and 1/4 & 2/4 & 3/4, respectively. deviations were calculated as the sum of square roots of absolute differences to the closest matched peaks of expected peak distributions. Peak estimates are shown in figure S6 and deviations between coverage and frequency based somy estimates in figure S8.
